# Impacts of wastewater treatment and the exposome on antibiotic resistance gene dynamics: Insights from Guadeloupe, a French Caribbean Island

**DOI:** 10.1101/2025.05.31.657145

**Authors:** Maria Alexa, Aleksandra Kovacevic, Mélanie Pimenta, Degrâce Batantou Mabandza, Ismaïl Ahmed, Sébastien Breurec, Christophe Dagot, Bich-Tram Huynh, Lulla Opatowski

## Abstract

Wastewater treatment plants (WWTPs) are global hotspots for the dissemination of antibiotic resistance genes (ARGs), primarily driven by different anthropogenic activities. Chemical compounds commonly found in wastewater, including antibiotics, biocides, non-steroidal anti-inflammatory drugs (NSAIDs), and heavy metals, along with environmental factors such as temperature and precipitation, might enhance the selection of antibiotic-resistant bacteria (ARB) even at low concentrations, inadvertently promoting the proliferation of ARGs. Since WWTPs can vary in their removal efficiency and influent compositions, it is important to assess their impact in influencing the relative abundance of ARGs and pollutant concentrations in discharged effluent. We analyzed data collected in Guadeloupe, French Caribbean, using random forest and lasso to investigate changes in the resistome and exposome within untreated wastewater influents and treated effluents from three WWTPs. These WWTPs represented three distinct wastewater continuums—hospital-based, urban non-touristic, and urban touristic—each with unique influent compositions. The study, which spanned four sampling campaigns from September 2021 to January 2023, revealed that the reduction in ARG abundance by the WWTPs was lower than expected. Of the 16 clinically relevant genes and mobile genetic elements examined, the relative abundance of *aph(3’)-III, bla*_OXA_, *bla*_SHV_, *bla*_TEM_, *ermB, intI1, qnrS*, and *tetM* was reduced by 59.8% to 89.9% across the three WWTPs. An increasing trend in the relative abundance of the *mcr-1* gene was observed, showing a 9.52-fold increase in the treated effluent in the hospital continuum. Of the nine antibiotics tested, trimethoprim levels dropped by 70.8% in the touristic continuum and 78.8% in the non-touristic, while ciprofloxacin saw a significant 64.4% reduction only in the touristic continuum, while no other antibiotics or biocides among the eight tested were effectively removed. Regarding heavy metals, significant reductions were observed only in the non-touristic continuum, with cadmium (Cd), copper (Cu), and mercury (Hg) concentrations reduced by 32.5%, 21.1%, and 36.2%, respectively. Our findings provide insights into the associations between environmental factors and the relative abundance of ARGs in Guadeloupe. Generally, depending on the continuum, biocides, the antibiotic erythromycin, certain heavy metals (As, Cd, Cu, Cr, and Hg) and water temperature were associated with the relative abundance of clinically relevant ARGs. This study was not able to assess the impact of climate. Future research should include longitudinal studies with more frequent sampling to better investigate the evolution of antibiotic resistance in bacterial species and its associated drivers, including pollutants and climate-related factors.

## 1. Introduction

Antimicrobial resistance (AMR) poses a significant global threat to human health and is projected to become the leading cause of death by 2050, representing a critical public health issue on a worldwide scale (Hou et al., 2023; Kwon and Powderly, 2021). In 2019, it was estimated that antibiotic-resistant pathogens were associated with 4.95 million deaths, with 1.27 million of these deaths being directly attributable to antibiotic resistance (Murray et al., 2022). Antibiotic-resistant bacteria (ARB) and antibiotic resistance genes (ARGs) have been detected in various habitats, including soil, water, wildlife and human-impacted areas. Anthropogenic activities, particularly the clinical use of antibiotics, are the main drivers of their dissemination (Bougnom and Piddock, 2017; Henriot et al., 2024; Larsson and Flach, 2022; Zhang et al., 2022).

Water plays a crucial ecological and evolutionary function within the context of AMR, facilitating the interaction of bacteria that originates from diverse habitats. This creates an ideal setting for ARGs exchange, mainly by horizontal gene transfer, among various bacterial species. ARGs are frequently present in various water sources, typically due to contamination from human activities. Wastewater from urban areas, agricultural and farming practices, and hospitals is a significant anthropogenic source of AMR in water environments (Bougnom and Piddock, 2017; Fouz et al., 2020; Reddy et al., 2022; Sambaza and Naicker, 2023). It contains high levels of nutrients and bacteria, along with numerous residual biological and chemical pollutants. Due to the clinical use of antibiotics, hospital effluent water is a major source of antibiotic compounds and has a higher concentration of ARB, ARGs, and antibiotic residues compared to other wastewater sources (Buelow et al., 2020; Zhu et al., 2023). The impact of anthropogenic activities on the environment has been recognized as a critical knowledge gap in understanding the evolution of antibiotic resistance (Larsson et al., 2018).

Resistome, a term first proposed by D’Costa and the colleagues refers to the collection of all ARGs present in a specific environment, organism, or microbiome (D’Costa et al., 2006). Wastewater contains antibiotics, disinfectants, and metals that, even at low concentrations, can create a selection pressure for antibiotic resistance. In the wastewater context, the resistome refers to ARGs and mobile genetic elements (MGEs) present in both influent and effluent. These elements confer resistance to antibiotics such as beta-lactams, macrolides, quinolones, sulfonamides, tetracyclines and can be carried by a wide range of bacteria.

The exposome, a term that was first introduced in human health epidemiology to describe “the totality of human environmental exposures” (Wild, 2005), in this context refers specifically to chemical compounds commonly found in wastewater from hospitals, urban areas, agricultural lands, and farms. These chemical compounds – including antibiotics, biocides, and heavy metals - along with other factors such as temperature and precipitation, might enhance the selection of ARB in wastewater, even in low concentrations (Harris et al., 2013; Sambaza and Naicker, 2023).

Understanding how the exposome affects the resistome and how pollution by selective agents may drive the evolution of resistance is essential to act on the reduction of AMR. Overlooking this critical factor could have serious health consequences, particularly in low- and middle-income countries where inadequate infrastructure for managing human and animal waste leads to increased environmental emissions of both resistant fecal bacteria and residual antibiotics (Kookana et al., 2014; Larsson and Flach, 2022).

Wastewater treatment plants (WWTPs) act as a link between human activities and the environment. Their primary purpose is to remove organic matter and nutrients, such as phosphorus and nitrogen, from urban wastewater before discharging the treated water into surface water bodies, such as rivers and oceans. However, despite classical treatment processes that result in around 2-log (99%) reduction in bacterial load, WWTPs are not designed to specifically eliminate ARBs and ARGs. Additionally, antibiotics often present in WWTPs in sub-inhibitory concentrations can promote transfer and selection of ARGs, mainly by horizontal gene transfer, contributing to the spread of AMR (Berendonk et al., 2015; Bruchmann et al., 2013; Harris et al., 2013; Partridge et al., 2018; Sambaza and Naicker, 2023; Stanton et al., 2022). This selective pressure enables ARBs to thrive and persist within the treatment environment, turning WWTPs into drivers of AMR (Berglund et al., 2023; Karkman et al., 2018; Manaia et al., 2018; Rizzo et al., 2013).

Last year, the Council and the European Parliament reached an agreement on the revised Urban Wastewater Treatment Directive. This new legislation updates the EU’s water management and urban wastewater treatment standards. It introduces several key measures, including a requirement to monitor the presence of AMR in urban wastewater, at upstream and downstream of WWTPs in cities with more than 100,000 population equivalent, as well as monitoring and removal of micropollutants. These measures aim to protect the environment from being adversely affected by insufficiently treated urban wastewater discharges and to contribute to the protection of public health in accordance with the One Health approach. They also incorporate additional water treatment applications, such as secondary treatment (i.e., the removal of biodegradable organic matter), before the water is discharged into the environment (European Parliament, 2024). Given that WWTPs can vary in their removal efficiency and influent compositions, it is important to assess their impact the relative abundance of ARGs and pollutant concentrations discharged into the effluent aquatic environment.

In this study, we investigate the impact of WWTPs on the relative abundance of 16 clinically relevant ARGs and MGEs in wastewater, as well as on the concentrations of chemical compounds in the surrounding exposome. We analyze and compare three distinct wastewater pathways: hospital-based, urban non-touristic, and urban touristic in Guadeloupe, a French Caribbean Island, each with varying influent wastewater compositions. Our focus is on identifying the resistome and exposome variables that are most affected by wastewater treatment processes and the exposome factors associated with relative abundance of antibiotic resistance genes.

## 2. Materials and methods

### 2.1 Sampling locations and study design

Guadeloupe, a French overseas island, has a population size of 384,315 inhabitants (2021) with high touristic activity (960,000 tourists per year in 2022). Sampling was conducted in three distinct regions of the island to capture and represent the diverse wastewater continuums shaped by various anthropogenic and tourism-related activities. These factors influence the characteristics of each wastewater pathway: (A) hospital continuum (wastewater from the University Hospital Center of Guadeloupe combined with urban wastewater from three major cities: Pointe-à-Pitre, Les Abymes, and Baie-Mahault), (B) non-touristic continuum (urban wastewater from a small town with minimal tourism activity, Lamentin, which is surrounded by poultry farms), and (C) touristic continuum (urban wastewater from Le Gosier, a small town with high tourism activity, including wastewater from hotels and a dialysis-specialized clinic) (Figure 1).

**Figure 1.**
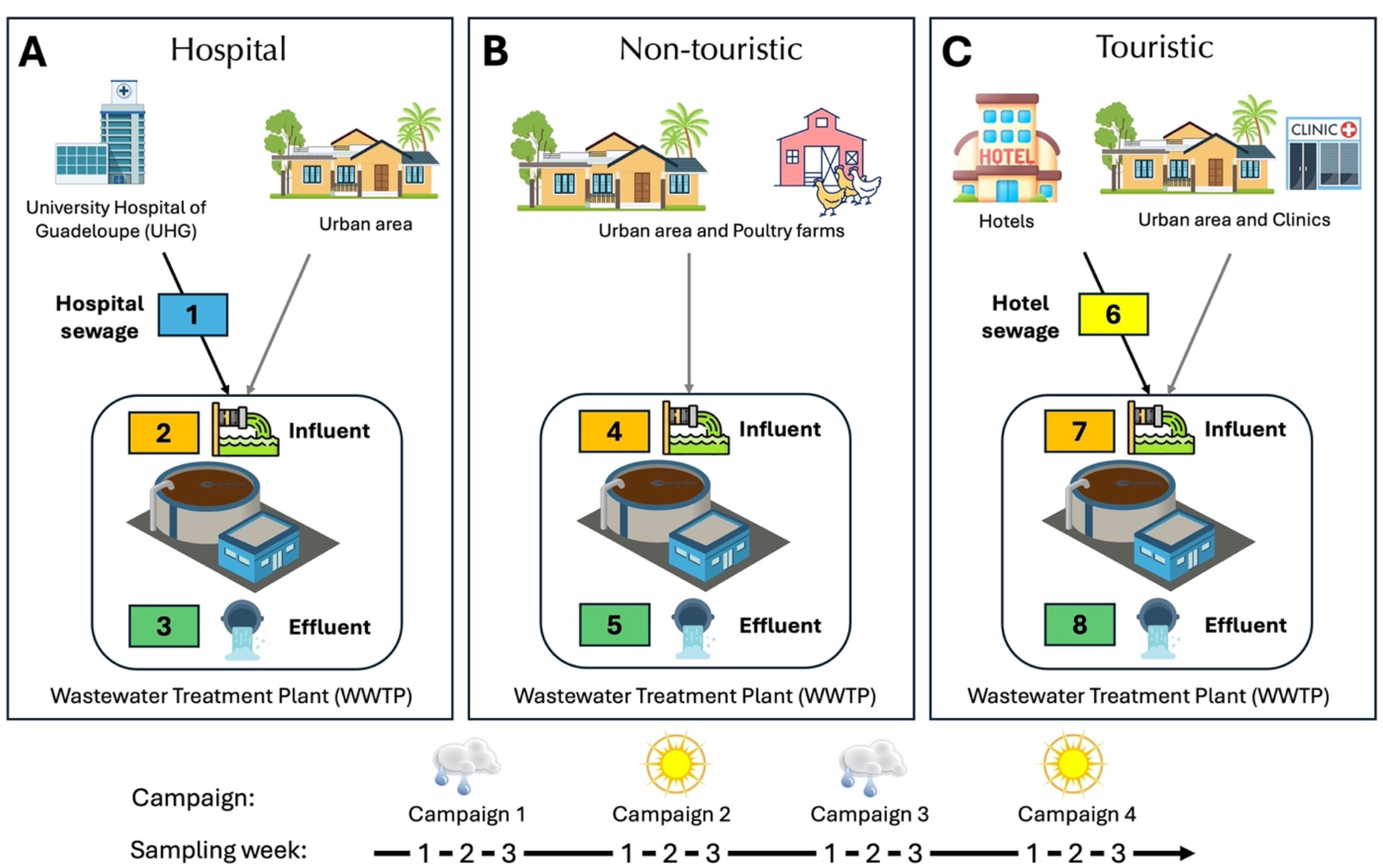
Wastewater samples were collected from three wastewater pathways in Guadeloupe, representing **(A)** Hospital, **(B)** Non-touristic, and **(C)** Touristic wastewater continuums. During each sampling campaign, samples were collected once a week for three consecutive weeks across eight sampling points. For a detailed description of sampling points 1-8, see Table S1 in the Supplementary Materials.

Wastewater samples were collected across four successive sampling campaigns, comprising two dry seasons and two rainy seasons. Over the course of each campaign, samples were collected once a week for three consecutive weeks from influent and effluent catch points along each wastewater continuum. A total of 96 wastewater samples, each measuring 2 liters, were collected throughout the 12-week sampling period. For further details on the sampling sites, refer to Table S1 in the Supplementary Materials. Sampling campaigns for the touristic and non-touristic continuums were conducted during rainy and dry seasons in September/October 2021 (rainy), January/February 2022 (dry), July/September 2022 (rainy), and January/February 2023 (dry). Hospital wastewater continuum had only three sampling campaigns conducted in January 2022 (dry season), September 2022 (rainy season), and January 2023 (dry season). The first sampling campaign from the hospital continuum was excluded from the analyses due to the low quality of the initial sample collection. This campaign did not accurately represent the rainy season in Guadeloupe, as it was conducted during a period with no rainfall in July, despite being expected to occur during this time.

### 2.2 Sample processing and analysis

Upon transportation to the laboratory maintaining water samples at 4°C, samples were filtered within 24h using a 0.22 μm filter (Millipore, Dominique Dutscher, Paris, France). The volume of water filtered varied according to the turbidity of the sample until the filter was saturated and was carefully recorded for each sample. Each filter was stored at −80°C in empty haemolysis tubes and then shipped on dry ice to Limoges for DNA extraction. DNA from each filter was extracted using the DNeasy PowerWater kit (Qiagen®) according to the manufacturer’s instructions. The DNA concentration was measured by Qubit™ dsDNA Broad Range Assay Kits (Invitrogen™). DNA was then diluted or concentrated to a final concentration of 10 ng/μL for downstream analysis. DNA was stored at −20°C until its use.

The relative abundance of genes, including 80 targeted genes that confer resistance to antibiotics categorized into 16 resistance gene classes, and quaternary ammonium compounds (QACs), was measured using high-throughput qPCR. Nine MGEs and three integron integrase genes (*intI1, intI2*, and *intI3*) were also targeted, as described previously (Buelow et al., 2018, 2017) (Table S2, Supplementary Materials). The bacterial 16S rRNA gene was targeted using universal primers. Real-time PCR was carried out using the 96.96 BioMarkTM Dynamic Array for Real-Time PCR (Fluidigm Corporation) (Buelow et al., 2018, 2017). PCRs were performed in triplicate (Figure S1, Supplementary Materials). The positive control was a wastewater sample that included every targeted gene, while the negative control was a sterile DNA-free water. The focus of the analysis was on 16 clinically-relevant ARGs and MGEs that confer resistance to aminoglycosides (*aac(6’)-lb, aph(3’)-III*), beta-lactams (*bla*_CTX-M_, *bla*_KPC_, *bla*_NDM_, *bla*_OXA_, *bla*_SHV_, *bla*_TEM_, *bla*_VIM_), macrolides (*ermB*), methicillin (*mecA*), polymyxin (*mcr-1*), quinolones (*qnrS*), sulfonamides (*sul1*), and tetracyclines (*tetM*), including a class I integron (*intI1*).

Normalized abundance of ARGs and MGEs was defined using the abundance of the 16S ribosomal RNA (rRNA) gene (2^(-(CT_ARG_-CT_16S rRNA_)). A log2 transformation was applied to the ratio of the abundance of antibiotic resistance genes (CT_ARG_) to the abundance of the 16S rRNA gene (CT_16S rRNA_), where CT represents cycle threshold (Buelow et al., 2017). The normalized abundance was treated here as a concentration.

Nine antibiotics (azithromycin, ciprofloxacin, erythromycin, ofloxacin, pyrimethamine, sulfaguanidine, sulfamethoxazole, sulfathiazole, trimethoprim), eight biocides (albendazole, flubendazole, levamisole, pyrantel, pyrimethamine, sulfoxaflor, tebuconazole, thiabendazole) and three NSAIDs (diclofenac, ibuprofen, ketoprofen) were measured after solid-phase extraction (SPE) by liquid chromatography coupled with tandem mass spectrometry (LC-MS/MS). For SPE, samples were loaded into Oasis HLB 3cc (60 mg) cartridges from Waters. The final dry residues were then reconstituted and injected into the LC-MS system. To conduct the quantitative analysis of the compounds, a multiple reaction monitoring (MRM) mode on a 5500 QTRAP (Sciex) mass spectrometer was used.

The concentrations of heavy metals in the water samples were measured using inductively coupled plasma mass spectrometry (ICP-MS) with the ICP-MS NexIon 350D (Perkin Elmer). Scandium (Sc), yttrium (Y), indium (In), and rhenium (Re) were used as internal standards. Metadata containing information on physicochemical properties and climate parameters, including temperature and precipitation, were collected during each sampling campaign (Table S2, Supplementary Materials).

### 2.3 Evolution of gene abundance and the pollutant concentration after the WWTP

A quality control filter step was initially applied to assess qPCR replicate variability, as detailed in sections 1.3 and 1.4 of the Supplementary Materials. This filtering step was designed to assess variability both within and between filters. Any relative abundance values that fell below the detection limit were replaced with a very low value of 1×10^−08^ (Srathongneam et al., 2024; Weiss et al., 2017). The first step in the analysis was to assess differences in the relative abundance (gene copies per 16S rRNA gene copy) of clinically relevant genes before and after the WWTPs. We also quantified the difference in concentration between influent and effluent samples for antibiotics, biocides, NSAIDs, and heavy metals. The analyses were assessed using the Wilcoxon signed-rank test for paired samples where p-values were adjusted using the Benjamini-Hochberg method, which controls the false discovery rate (Wilcoxon, 1945).

### 2.4 Variables in the resistome and exposome most impacted by WWTPs

The next step of the analysis was to identify the variables among the resistome and exposome the most shaped by the wastewater treatment. These variables are also the ones that characterize and predict the likely source of wastewater samples (influent vs. effluent) for each wastewater continuum. To identify these variables, we used the Random Forest Algorithm (RFA). Predictions of the most likely wastewater sampling source were based on measurements of the relative abundance of ARGs and MGEs as well as the concentration of pollutants, which served as predictor variables. We ran the RFA separately on the resistome and exposome datasets. For further details on the RFA pipeline please and ranking of our predictors refer to sections 1.1, 1.2, and Figure S4 in the Supplementary Materials.

### 2.5 Factors associated with relative abundance of antibiotic resistance genes

Exposome variables were factors analyzed in relation to the relative abundance of 16 clinically relevant ARGs and MGEs at specific wastewater sampling point (either influent – sampling points 2, 4 and 7 or effluent – sampling points 3, 5, 8 in Figure 1) within each wastewater continuum. To accomplish this, we employed the Least Absolute Shrinkage and Selection Operator (Lasso) technique (Tibshirani, 1996), minimizing the Bayesian Information Criterion (BIC) (Courtois et al., 2021). Each factor had either a positive (+) or negative (-) association with antibiotic resistance status depending on the sign of the coefficients. Lasso is a linear regression algorithm that performs L1 regularization, which adds a penalty technique by shrinking some coefficients towards zero, and a good method to fit models when data show high dimensionality.

Due to the high correlation among all heavy metals, we used Principal Component Analysis (PCA) to address multicollinearity and kept only the first two principal components (Figures S5-S7, Supplementary Materials). For other highly correlated explicative variables from the exposome, physicochemical properties, and climate parameters (threshold was set at >0.708) only one variable between the two was included in the model based on clinical/biological relevance. The significance threshold for this correlation analysis was set at an alpha level of 0.01 for Pearson correlations. With a sample size of n=12, the degrees of freedom are n-2, resulting in 10. Therefore, the threshold for a significant correlation at an alpha level of 0.01 was 0.708. Gene *bla*_NDM_ had too many data points below the detection limit and was consequently excluded from the Lasso analysis.

Let *P* represent the total number of covariates, *P*_*e*_ the number or exposome covariates, *P*_*pc*_ the number of physicochemical covariates, and *P*_*c*_ the number of climate covariates. Thus, *P = P*_*e*_ + *P*_*pc*_ + *P*_*c*_ + 1, where the additional 1 represents the covariate for the relative abundance of the upstream gene.

Let *X* denote the *N* × *P* matrix of covariates, defined as the concentration of the following vectors:

*x*_*u*_: the relative gene abundance found at the upstream sampling location.

*X*_*e*_: exposome (pollutant concentration) covariates of size *N* × *P*_*e*_, where *x*_*e, i*_ corresponds to the *i*th covariate.

*X*_*pc*_: physicochemical covariates of size *N* × *P*_*pc*_, where *x*_*pc, k*_ corresponds to the *k*th covariate.

*X*_*c*_: climate covariates of size *N* × *P*_*c*_, where *x*_*c, l*_ corresponds to the *l*th covariate.

Let *Y* be the *N*-vector of relative gene abundance found at the influent/effluent sampling location. For each influent and effluent sampling point, we have the following model for a given time *t*:

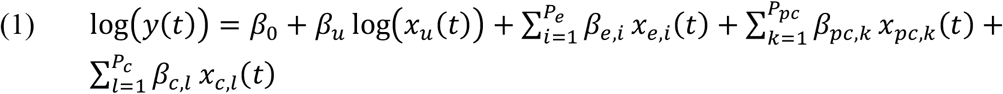

where *β*_0_ is the intercept, and *β*_*u*_, *β*_*e, i*_, *β*_*pc, k*_, *β*_*c,l*_ are the regression coefficients associated with the upstream gene covariate, exposome covariates, physicochemical covariates, and climate covariates, respectively.

The Lasso penalization term is defined as:

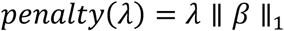

Meaning:

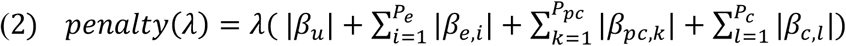

Thus, the penalized Lasso consists of estimating:

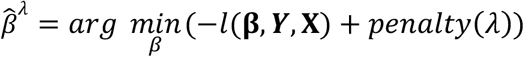

with *l the log* − *likelihood of the model* (1).

The BIC was implemented and defined for each *λ* as follows:

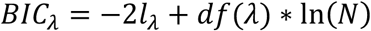

with N representing the number of observations, *l*_*λ*_ the log-likelihood, and *df*(*λ*) the effective degrees of freedom.

## 3 Results

### 3.1 Evolution of ARGs and MGEs after wastewater treatment

Among the 16 genes of interest, the relative abundance of *aph(3’)-III, bla*_OXA_, *bla*_SHV_, *bla*_TEM_, *ermB, intI1, qnrS*, and *tetM* was systematically reduced after wastewater treatment in all three continuums (Figure 2), with the average reductions ranging from 59.8% to 89.9% (Table S3, Supplementary Materials). Conversely, the relative abundance of *mcr-1* gene, which confers resistance to colistin via plasmids did not display a significant reduction after wastewater treatment in any of the three continuums. In fact, the mean relative abundance of *mcr-1* gene was 9.52-fold higher in treated effluent than in untreated influent wastewater in the hospital continuum (Table S3, Supplementary Materials).

**Figure 2.**
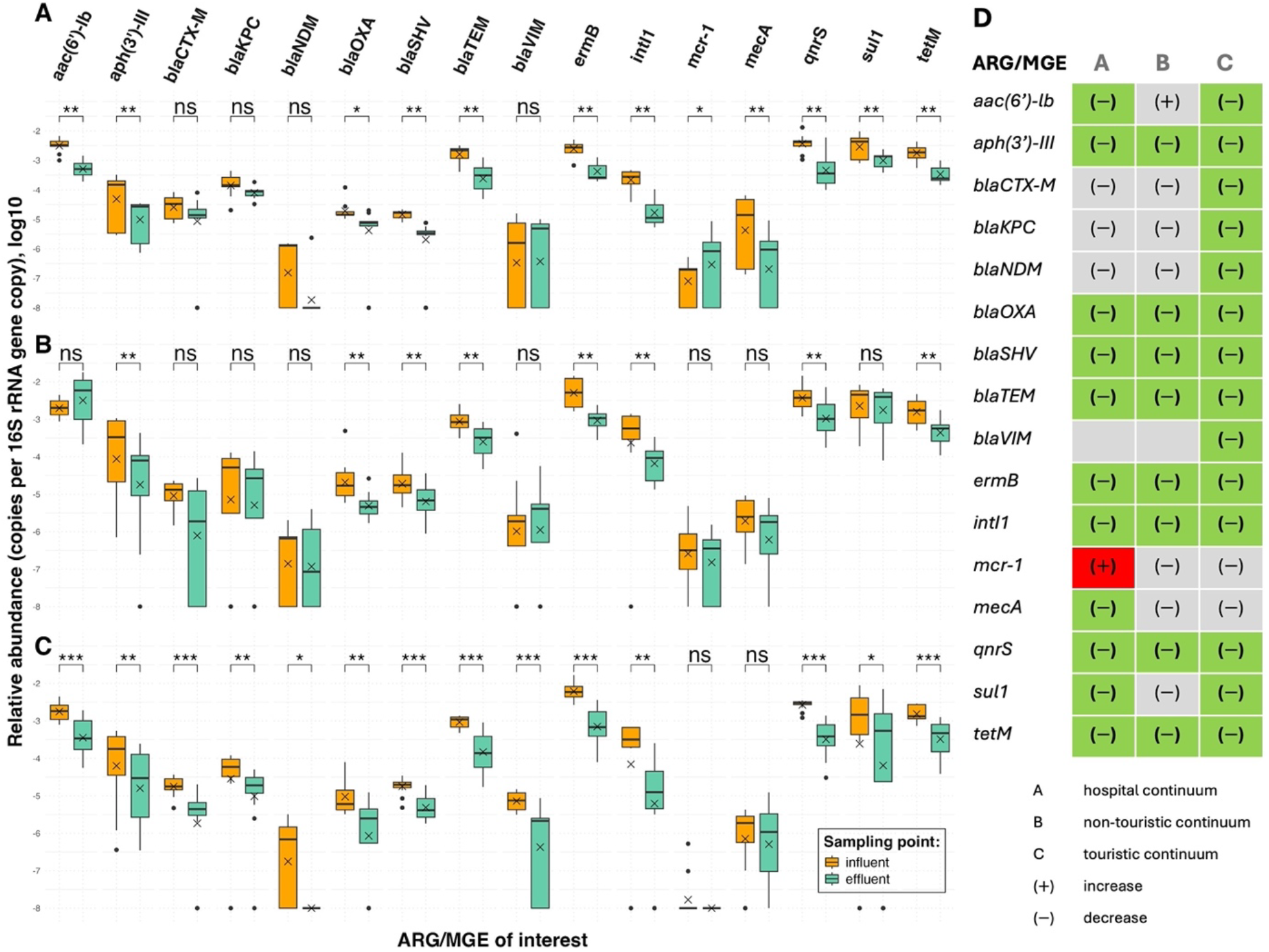
Relative abundance of 16 ARGs and MGEs of interest before (influent) and after (effluent) wastewater treatment. **(A)** Hospital, (**B)** Non-touristic, and (**C)** Touristic wastewater continuums. Statistical significance was assessed using the non-parametric Wilcoxon signed-rank test for paired samples where p-values were adjusted using the Benjamini-Hochberg method (Table S4). Barplots show medians while ‘x’ markers indicate mean values. (**D)** Summary table of changes in relative abundance of 16 genes of interest before (influent) and after (effluent) wastewater treatment across three wastewater continuums: significant increase (red), significant decrease (green) or no statistically significant change (grey). Table S3 shows % change in the average relative abundance from influent to effluent wastewater samples.

Across all sampling campaigns combined, the relative abundances of the genes *bla*_CTX-M_, *bla*_KPC_, *bla*_NDM_, and *bla*_VIM_ conferring resistance to beta-lactams were significantly reduced by WWTPs only in the effluent of the touristic wastewater continuum, with reductions ranging from 64.5% to 98.9% (Table S3, Supplementary Materials). Genes *aac(6’)-lb, bla*_NDM_ and *mecA* had higher relative abundance in the effluent of non-touristic and touristic continuums, but these changes were not statistically significant (Table S3, Supplementary Materials). Furthermore, the *ermB* gene, which confers resistance to macrolides, was by far the most abundant in the effluent water of both urban touristic and non-touristic continuums. In the hospital continuum, the *ermB* gene was among the top five in terms of relative abundance with *qnrS* and *sul1* genes having the highest relative abundances in this continuum (Table S3, Supplementary Materials).

### 3.2 Evolution of antibiotic concentrations after wastewater treatment

Across all three wastewater continuums, the concentration of antibiotics in the effluent wastewater samples generally showed no significant change compared to influent levels. The effluent concentration of trimethoprim (TMP) was reduced by 70.8% and 78.8% in the touristic and non-touristic continuums, respectively, while the concentration of ciprofloxacin (CIP) was significantly reduced only in the touristic continuum by 64.4%. All other antibiotics remained at similar concentration levels in the effluent wastewater as they were in the influent (Figure S8 and Tables S3 and S4, Supplementary Materials).

### 3.3 Evolution of biocide, non-steroidal anti-inflammatory drug, and heavy metals concentrations after wastewater treatment

The concentration of the biocides did not significantly change between influent and effluent wastewater samples (Figure S9, Supplementary Materials). Similarly, concentrations of NSAIDs did not significantly differ from the influent to effluent wastewater either, except for a decrease in ketoprofen concentration by 90.3% observed in the hospital wastewater continuum (Figure S10 and Table S3, Supplementary Materials). For heavy metals, concentrations of cadmium (Cd), copper (Cu), and mercury (Hg) were significantly reduced only by the non-touristic WWTP, with reductions of 32.5%, 21.1%, and 36.2%, respectively. However, the concentrations of arsenic (As), chromium (Cr), gadolinium (Gd), lead (Pb), and zinc (Zn) remained unchanged after wastewater treatment across all three continuums (Figure S11, Supplementary Materials). See Tables S3 and S4 in Supplementary Materials for more details.

### 3.4 Variables in the resistome and exposome most impacted by WWTPs

The Random Forest Analysis provided the top predictors that exhibited the most significant differences in abundance or concentration between wastewater sampling sources before and after wastewater treatment, therefore indicating the variables in resistome and exposome possibly most affected by WWTPs. Top variables are shown in Figure 3 and mean accuracy in Table 1. The main findings suggest that ARGs, including genes of interest, are significantly impacted by WWTPs. More specifically, in the effluent wastewater in all three continuums, there was reduced relative abundance of the *ermB* gene, which is associated with resistance to macrolides. The *ermB* gene was also among the top fifteen ARGs most affected by wastewater treatment in both the hospital and touristic continuums. Additionally, we found that targeted efflux pump genes (*acrA, mdtO, tolC, mdtL*, and *mdtF*) are among the top fifteen variables whose relative abundance is most reduced by WWTP C in the touristic continuum. Ciprofloxacin and copper were among the top exposome variables distinguishing influent and effluent sampling points in both the touristic and non-touristic wastewater continuums. Additionally, trimethoprim was a key differentiating factor in the non-touristic continuum (Figure 3). Antibiotics, including ciprofloxacin, azithromycin, erythromycin and trimethoprim, and the NSAID ketoprofen, were among the top exposome variables distinguishing influent and effluent sampling points in the hospital continuum (Figure 3).

**Table 1.** Random Forest results using either resistome or exposome data. Mean accuracy for all three continuums. Poor mean accuracy is highlighted in red.

|  | Input data |  |
| --- | --- | --- |
|  | Resistome | Exposome |
|  | Mean Accuracy (SD) | Mean Accuracy (SD) |
| Hospital | 91.3 (14.7) | 91.3 (12.2) |
| Non-touristic | 87.5 (13.1) | 80 (15.9) |
| Touristic | 94.2 (8.2) | 69.2 (18.2) |

**Figure 3.**
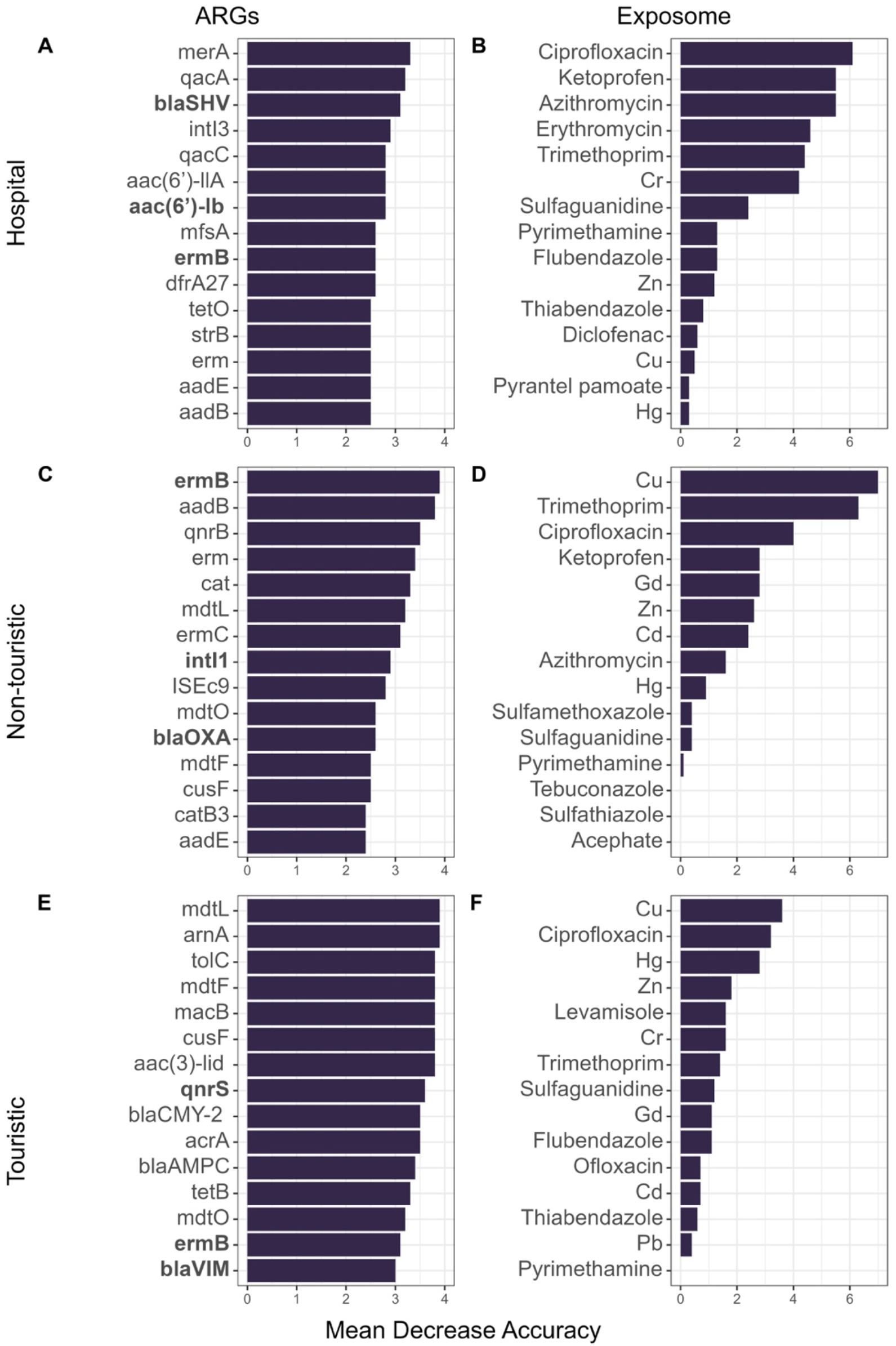
Ranking of top 15 mean decrease in accuracy (MDA) values in random forest analysis (RFA) using either resistome or exposome data, a statistical classification that indicates the importance of each variable. Clinically relevant genes are indicated in bold and confer resistance to aminoglycosides (*aac(6’)-lb*), beta-lactams (*bla*_OXA_, *bla*_SHV_, *bla*_VIM_), macrolides (*ermB*), and quinolones (*qnrS*) including a class I integron (*intI1*).

### 3.5 Factors associated with relative abundance of antibiotic resistance genes

In the final step of the analysis, we investigated key environmental drivers and anthropogenic factors associated with the relative abundance of resistance genes in the influent and effluent wastewater. These hypothesized factors included antibiotics, biocides, NSAIDs, heavy metals, physicochemical properties, and climate parameters. We also considered the relative gene abundance at points prior to the influent and effluent sampling locations. Specifically, the term “Upstream Gene” refers to the relative abundance of genes at sampling points 1 and 6 for the influent (not available for the non-touristic continuum) and at sampling points 2, 4, and 7 for the effluent sampling location.

For the influent sampling locations, the lasso analysis revealed that the relative abundance of ARGs in the wastewater coming directly either from the hospital or from hotels (“Upstream Gene”) exhibited a positive (+) association with the relative abundance of genes *aac(6’)-lb, aph(3’)-III, bla*_KPC_, *bla*_OXA_, *intI1, mcr-1, sul1*, and *tetM* found in the influent of their respective WWTP (Figure 4A and Figure 4E).

**Figure 4.**
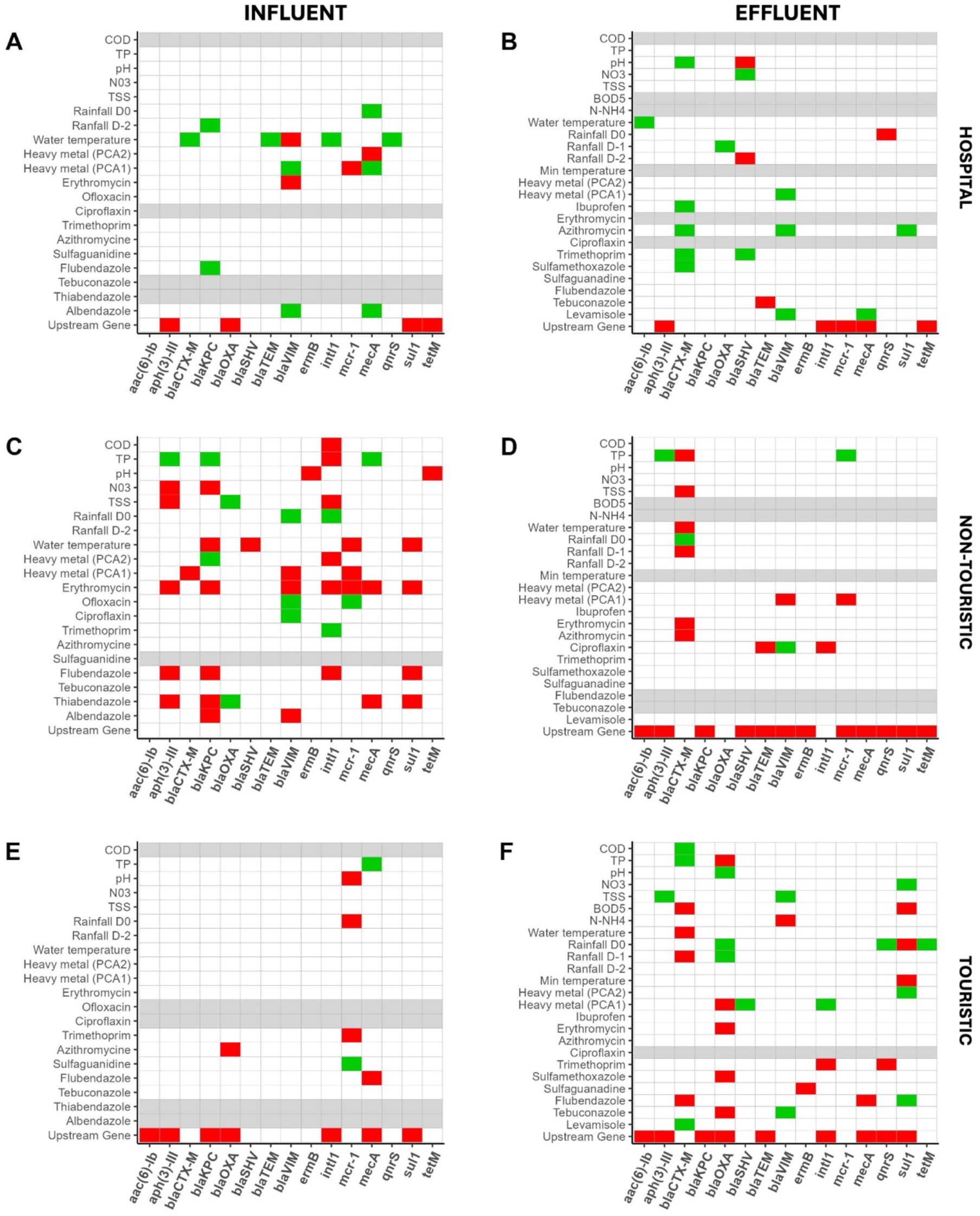
Heatmaps representing factors associated with relative abundance of genes of interest in the influent and effluent wastewater samples in hospital, non-touristic, and touristic wastewater continuums. Red color indicates a positive (+) association, green indicates a negative (-) association, white squares represent variables not selected by the lasso (null coefficient), and grey color represents variables removed due to correlation with another variable. The gene *bla*_NDM_ has been excluded from the analyses (COD = chemical oxygen demand; TP = total Phosphorus; TSS = total suspended solids; BOD = biochemical oxygen demand).

In general, most biocides (except for thiabendazole in the case of the gene *bla*_OXA_), the antibiotic erythromycin, some selected heavy metals (As, Cd, Cu, Cr, and Hg) and water temperature were positively (+) associated with the relative abundance of ARGs in the influent of the non-touristic continuum (Figure 4C). Conversely, this positive association was not widely observed in influent of the hospital and touristic continuums. Interestingly, in the hospital continuum (Figure 4A), water temperature showed a negative (-) association with four genes of interest, including an integron (*bla*_CTX-M_, *bla*_TEM1_, *qnrS*, and *intI1*), and a single positive association (+) with *bla*_VIM_.

For the effluent sampling locations, the factor Upstream Gene had a positive (+) association with the relative abundance of most ARGs in the effluent water across both non-touristic and touristic continuums (Figure 4D and Figure 4F). In the hospital continuum (Figure 4B), the antibiotic azithromycin had a negative (-) association with the *bla*_CTX-M_, *bla*_VIM_, and *sul1* genes. Additionally, pH and rainfall two days before sample collection in the hospital continuum had a positive (+) association with the relative abundance of *bla*_SHV_ gene.

## 4. Discussion

A total of 96 wastewater samples were analyzed over a 16-month period (September 2021-January 2023) from eight sites across three wastewater continuums in Guadeloupe. The results showed that the reduction in ARG abundance after wastewater treatment plants was lower than anticipated. While some antibiotics were reduced by the wastewater treatment plants, biocides were not effectively removed. Additionally, heavy metal removal was also significantly lower than the generally expected rates. We examined the key environmental drivers and anthropogenic factors among the exposome, physicochemical properties, and climate parameters associated with the relative abundance of ARGs before and after wastewater treatment. This study aimed to gain insights into how WWTPs and environments affected by different human activities contribute to the dissemination of ARGs (Samreen et al., 2021; Wales and Davies, 2015; Wang et al., 2020; Webber et al., 2015).

Among the 16 clinically relevant ARGs and MGEs studied, the relative abundance of *aph(3’)-III, bla*_OXA_, *bla*_SHV_, *bla*_TEM_, *ermB, intI1, qnrS*, and *tetM* was consistently reduced by 59.8% to 89.9% after wastewater treatment in all three continuums. However, the reduction in ARGs abundance observed was below the expected reduction of 2.0 ± 1.0 log and a removal efficiency of 90%-100%, which is typically reported for activated sludge and other treatment processes (Wang et al., 2020). The observed 9.52-fold increase in the relative abundance of *mcr-1* gene in treated effluent wastewater in the hospital continuum is an important and concerning observation, given that *mcr-1* is colistin resistance gene mediating resistance against last resort antibiotic polymyxin E.

According to a study conducted in Vietnam, the primary source of *mcr-1* gene in urban water environments is human feces (Nguyen et al., 2021). However, it appears that WWTPs do not significantly impact the survival and persistence of *mcr-1* gene carrying bacteria (Hembach and Schwartz, 2017). The resilience of certain ARGs, such as *mcr-1*, during treatment processes raises concerns about the potential for these genes to persist and spread, highlighting the importance of monitoring and optimizing WWTP operations to address such challenges (Marti et al., 2013). Notably, the relative abundances of four genes of clinical interest conferring resistance to extended-spectrum beta-lactams (*bla*_CTX-M_) and carbapenems (*bla*_KPC_, *bla*_NDM_, *bla*_*VIM*_) were significantly reduced only by the WWTP in the touristic wastewater continuum, despite this plant not being compliant with regulatory standards. This is particularly interesting as resistance to beta-lactamases and carbapenems represents an important public health issue worldwide, limiting treatment options for severe human infections.

The concentration of the antibiotic trimethoprim (TMP) was significantly reduced by WWTPs in both the non-touristic and touristic continuums, with reductions of 70.8% and 78.8%, respectively. The antibiotic ciprofloxacin (CIP) was reduced by 64.4% only in the touristic continuum. Interestingly, this reduction was not observed for sulfamethoxazole (SMX), which is commonly administered in combination with trimethoprim as TMP-SMX. In contrast, SMX concentration levels increased in the non-touristic continuum, although this change was not statistically significant. The lack of reduction in SMX concentration after wastewater treatment may be attributed to its long half-life, which can exceed a year. Additionally, sulfamethoxazole is considered poorly biodegradable in activated sludge systems, commonly used in WWTPs. However, it can undergo degradation in anoxic or anaerobic conditions, where different microbial communities and redox conditions might facilitate its breakdown (Dagot, 2018). Therefore, the persistence of SMX in the environment raises concerns about the effectiveness of current wastewater treatment processes in removing this compound and highlights the need for improved treatment technologies that can address such pollutants.

Additionally, the antibiotics ofloxacin, pyrimethamine, and sulfaguanidine also showed an increasing trend in the non-touristic continuum associated with WWTP which met regulatory standards although this trend was not statistically significant. The half-lives of these antibiotics in wastewater can vary significantly based on conditions such as temperature, pH, and the presence of other compounds, but general estimates are that these antibiotics can last up to two weeks. This finding is consistent with recent review indicating that WWTP facilities often fail to adequately eliminate antibiotics, with removal rates ranging between 53% and 78% (Wang et al., 2020). In line with this, our findings reveal that no other antibiotics, except for TMP and CIP, were effectively removed by the WWTPs (Figure S5, Supplementary Materials).

The concentrations of eight biocides and three NSAIDs were largely unaffected by WWTP processes, except for the ketoprofen, which was reduced by 90.3% in the hospital continuum’s effluent wastewater. Most studies show very variable reductions and indicate that conventional wastewater treatment typically reduces diclofenac by 0% to 69% and ibuprofen by more than 90% (Dagot, 2018). Overall, NSAIDs removal efficiency in wastewater treatment plants has been reported to be around 4–89%, 8–100%, and 17–98% for diclofenac, ketoprofen, and ibuprofen, respectively (Luo et al., 2014; Mussa et al., 2022). Removal rate of biocides in WWTPs can vary depending on the biocide but is typically between 60% and 90%. Our findings are concerning, as we observed no change in the concentration of any of the eight biocides before and after wastewater treatment.

The removal rate of heavy metals in water by WWTPs can vary widely depending on the specific metal, the treatment processes used, and the characteristics of the influent water. In our study, we observed that only the non-touristic WWTP significantly reduced the concentrations of heavy metals. Specifically, Cd, Cu, and Hg were reduced by 32.5%, 21.1%, and 36.2% in the effluent water, respectively. Generally, removal rates for these metals in the soluble phase are expected to range from 60% to 95% suggesting that heavy metal removal in the studied WWTPs is quite low.

The *ermB* gene, which confers resistance to macrolides, was the most affected by WWTP in the non-touristic continuum. Additionally, all five targeted efflux pump genes (*acrA, mdtO, tolC, mdtL*, and *mdtF*) were among the top ARGs whose relative abundance was most reduced by WWTP in the touristic continuum (Figure 3E). The antibiotic ciprofloxacin (CIP) and copper (Cu) were among the top variables in the exposome most reduced by the wastewater treatment process across all three continuums (Figure 3). These variables help characterize the most likely wastewater sampling source (influent vs. effluent).

The relative abundance of ARGs found upstream (“Upstream Gene”) was identified as one of the main factors positively associated with the relative abundance of genes of interest in both influent and effluent wastewater. In the non-touristic continuum, biocides, the antibiotic erythromycin, and selected heavy metals (As, Cd, Cu, Cr, and Hg) were also positively associated with the relative abundance of genes of interest in the influent water. This positive association was not widely observed in the touristic and hospital continuums. Additionally, in the hospital continuum, biocides and azithromycin were generally negatively associated with the relative abundance of certain genes of interest in the effluent water.

In the influent water of the hospital continuum, water temperature showed a negative association with the relative abundance of four genes of interest (*bla*_CTX-M_, *bla*_TEM_, *qnrS, intI1*) and a positive association with the gene *bla*_VIM_. In contrast, this was not observed in the effluent water of the hospital continuum. While rainfall is generally associated with an increase in the relative abundance of ARGs (Di Cesare et al., 2017; Magnano San Lio et al., 2023), the negative association between water temperature and ARGs contrasts with findings from other studies. For instance, the gene *tetM* has been shown to positively associate with water temperature (Yu et al., 2023). Interestingly, at higher temperatures, the relative abundance of naturally occurring ARGs in river biofilms increases, although this is accompanied by a reduced success rate of invading foreign ARGs from wastewater in establishing within these communities (Bagra et al., 2024). In the hospital continuum, water temperature ranged between 25ºC and 29.5ºC across sampling weeks, suggesting that fluctuations within this range may significantly impact some genes of interest negatively.

In France, 90% of urban wastewater is treated in accordance with the Urban Wastewater Treatment Directive (UWWTD), surpassing the EU average of 76% (Water Information System for Europe). However, this national average masks significant regional disparities. In 2021, 78% of wastewater treatment systems in Guadeloupe with a nominal capacity of 2,000 population equivalents (PE) or more were non-compliant with official regulatory standards. Among the three wastewater continuums examined in this study (Figure 1), the hospital and touristic facilities were non-compliant, whereas the plant in the non-touristic continuum (Figure 1B) met the regulatory requirements set by the Department of the Environment, Planning and Housing (Ministère de la Transition Écologique, 2023). While this compliance status may have influenced concentrations of certain pollutants (e.g., heavy metals), it did not appear to impact AMR.

Our study has certain limitations. While the number of samples collected aligns with those in similar studies, we conducted four sampling campaigns, each consisting of three points, with each point taken during a different week. Although more frequent sampling, either throughout the year or within each campaign, might have increased the statistical power of our analysis, our primary goal was to capture both seasons present in Guadeloupe. Furthermore, some measurements of gene abundance fell below the detection threshold, potentially limiting the comprehensiveness of our data. Consequently, we cannot determine whether these genes were present at very low abundance or entirely absent from the wastewater samples.

Nevertheless, our results provide valuable insights into the associations between environmental factors and the relative abundance of ARGs. The geographical characteristics of territories such as Guadeloupe, including insularity, climate, rainfall, and tourism, present unique challenges for wastewater treatment. These challenges impact water resources, the broader environment, public health, and human activities. One major concern is the risk of surface water contamination in mangrove areas and tourist zones, often caused by accidental discharges or inadequately treated wastewater. Efforts must be made to improve treatment technologies and infrastructure, tailored to the specific needs of these territories. Furthermore, upcoming European regulations will strengthen standards and impose stricter requirements for monitoring wastewater discharges, creating additional challenges for maintaining treatment efficiency and ensuring regulatory compliance. Improving wastewater management in island environments will require specific measures focused on integrated water management and enhanced monitoring, while taking into account climatic hazards, infrastructure upgrades, economic development, and the protection of public health and the environment.

## 5. Conclusions

- WWTPs in Guadeloupe partially reduce the abundance of clinically relevant ARGs and associated pollutants, with variations depending on the type of the wastewater continuum.
- A concerning 9.52-fold increase in *mcr-1* (colistin resistance) was observed post-treatment in the hospital continuum, highlighting WWTP inefficiency in eliminating certain high-risk ARGs.
- A significant association was observed between specific ARGs and chemical pollutants.
- Future research extending such spatiotemporal designs based on longitudinal follow up with increased sampling frequency should be done to extend our understanding of the roles of pollutants, environmental conditions, and climate variability in shaping the antibiotic resistance patterns in water systems.

