## Supplementary Material for "Impacts of wastewater treatment and the exposome on antibiotic resistance gene dynamics: Insights from Guadeloupe, a French Caribbean Island"

### *Appendix 1*

**List of Acronyms and Abbreviations**

| | |
|---|---|
| WWTP | Wastewater Treatment Plant |
| *Antibiotics* | |
| AZ | Azithromycin |
| CIP | Ciprofloxacin |
| ERY | Erythromycin |
| OFX | Ofloxacin |
| PM | Pyrimethamine |
| SGN | Sulfaguanidine |
| SMX | Sulfamethoxazole |
| STZ | Sulfathiazole |
| TMP | Trimethoprim |
| *Biocides* | |
| ACE | Acephate |
| ABZ | Albendazole |
| FLU | Flubendazole |
| LEV | Levamisole |
| PYR | Pyrantel pamoate |
| SUL | Sulfoxaflor |
| TEB | Tebuconazole |
| TBZ | Thiabendazole |

#### 1.1 Random Forest Algorithm Pipeline

RFA pipeline included three steps: (1) data splitting, (2) grid search cross-validation, and (3) the final evaluation (step 1 in Figure S4). First, data was split into a training set and a testing set. Data splitting was done randomly with 75% of the data being considered as the training set and 25% of the data as the testing set. Given the significant time gap between the campaigns, spanning from September 2021 to January 2023, there may have been potential variations or discrepancies to consider. To mitigate the impact of data splitting on the resulting predictors and accuracy, we conducted multiple random forests (n=20) and averaged the results across multiple random data splits (Buelow et al., 2020). The second step of our pipeline was aimed at finding the best hyperparameters based on the analysis of the training set. Here, the parameter we wanted to tune was the *mtry* parameter defined as the number of randomly selected predictor variables out of which each split is selected when growing a tree (Probst et al., 2019). The default value for the *mtry* parameter is the square root of the number of variables. We created a grid for tuning our parameters ranging from 2 to the square root of the number of variables and used repeated 5-fold cross-validation. That means that our training dataset was randomly partitioned into five subsets (folds). The model was trained and evaluated five times, each time with a different random partitioning.  Each iteration assessed model accuracy for a specific *mtry* parameter value. This cross-validation process was repeated three times with distinct random splits. Ultimately, the value of the *mtry* parameter that yielded the highest accuracy was selected. The final model was then applied to the testing set using this optimal *mtry* parameter value. In the third step, we used the accuracy to evaluate the performance of our model. The accuracy, which is the probability that the model prediction is correct, has been suggested to be good metric for multi-class classification problems (Grandini et al., 2020). The models were run in R version 4.3.0, using the 'caret' package (Kuhn, 2008)

#### 1.2 Measure of predictor importance

The Random Forest Algorithm (RFA) pipeline is detailed in section 1.1. To rank our top predictors, i.e., the factors whose abundance or concentration is most affected by WWTP, we used the Mean Decrease Accuracy score (Chen and Ishwaran, 2012) (step 2 in Figure S4). The classification accuracy after permutation of a predictor variable's data is subtracted from the accuracy without permutation and averaged over all the trees in the random forest. Finally, the average over all the trees is divided by the standard deviation of the accuracy differences.

$$Mean\ Decrease\ Accuracy = \frac{Mean\ (Decrease\ in\ Accuracy\ of\ Trees)}{Standard\ Deviation\ (Decrease\ in\ Accuracy\ of\ Trees)}$$

#### 1.3 Statistical Framework for Assessing Intra- and Inter-Filter Variability

*Intra-Filter Variability*

To assess variability among the three technical replicates within each filter, we compared the distribution of CT values across all target genes. The underlying hypothesis was that replicates (Figure S1) should exhibit consistent distributions. Statistical analysis was performed using the Wilcoxon signed-rank test, with p-values adjusted using the Benjamini-Hochberg method and a significance threshold of 0.05. If significant differences were detected, one or all replicates for the affected filter were removed. Specifically, if a single replicate exhibited a statistically significant deviation from the others, only that replicate was removed. Conversely, if all replicates were statistically different from one another, they were all excluded. After this filtering step, the relative abundance of genes was calculated, and the mean value for each filter was retained for further analysis.

*Inter-Filter Variability*

The next step was to assess the consistency of measurements across different filters. We analyzed differences in the distribution of mean relative abundances for all target genes between filters using a Kruskal-Wallis test. Dunn's post-hoc test was then applied to identify specific pairwise differences between filters, with p-values adjusted using the Benjamini-Hochberg method and a significance threshold of 0.05.

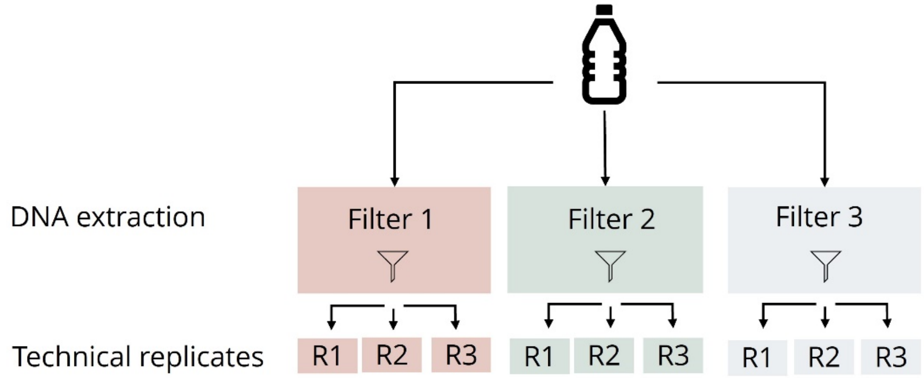


**Figure S1. Sample processing.** Each water sample was divided into three subsamples, each filtered using an identical filter. PCRs were performed in triplicate to obtain technical replicates. Here, for a given filter, R*i* denotes the *i*th replicate of that filter.

##### 1.4 Application of Statistical Filtering for Quality Control and Data Refinement

*Intra-Filter Variability*

No significant differences were observed between technical replicates within individual filters for both the hospital and non-touristic continuums (Figure S2). However, in the touristic continuum, statistical differences were detected between technical replicates in the following cases: Filter 3 during week 7 of the touristic sample, Filter 2 during week 4 of the influent sample, and Filters 2 and 3 during week 4 of the effluent sample.

Based on these results, the following technical replicates were removed:

- Touristic continuum, week 7 – Touristic hotels sample, Filter 3, Replicate 2
- Touristic continuum, week 4 – Influent sample, Filter 2, Replicate 1
- Touristic continuum, week 4 – Effluent sample, Filter 2, all replicates
- Touristic continuum, week 4 – Effluent sample, Filter 3, Replicate

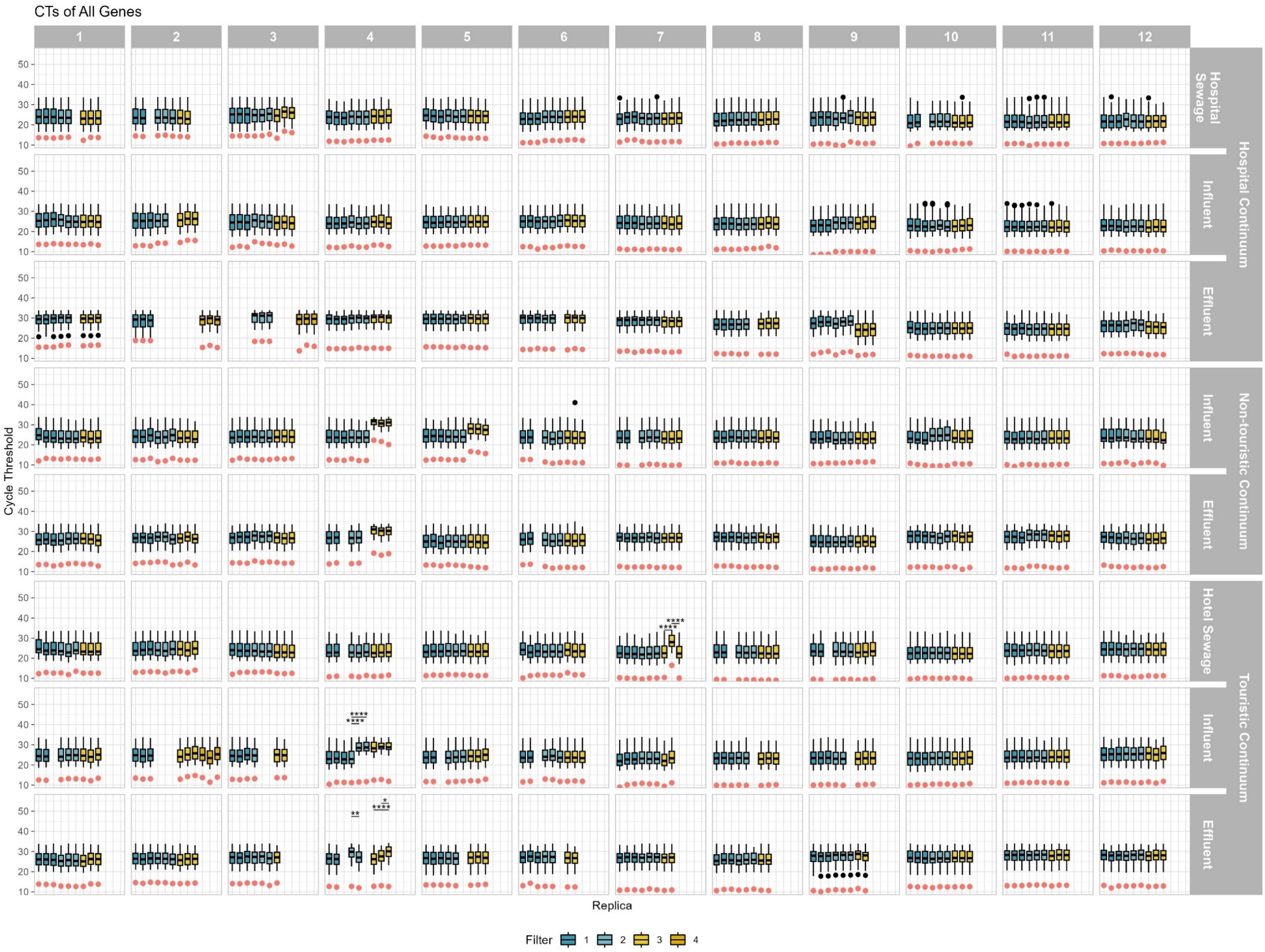

CTs of All Genes
1
2
3
4
5
6
7
8
9
10
11
12
Hospital Sewage
Influent
Effluent
Hospital Continuum
Influent
Effluent
Non-touristic Continuum
Hotel Sewage
Influent
Effluent
Touristic Continuum
Cycle Threshold
10
20
30
40
50
Replica
Filter 1 2 3 4

**Figure S2. Cycle threshold distribution of all ARGSs and MGEs before (influent) and after (effluent) wastewater treatment for each technical replicate.** The horizontal numbers represent the sampling weeks. Red dots indicate the cycle threshold of the 16S gene. Asterisks indicate significant differences between technical replicates (* p<0.05, ** p<0.01, *** p<0.001, **** p<0.0001).

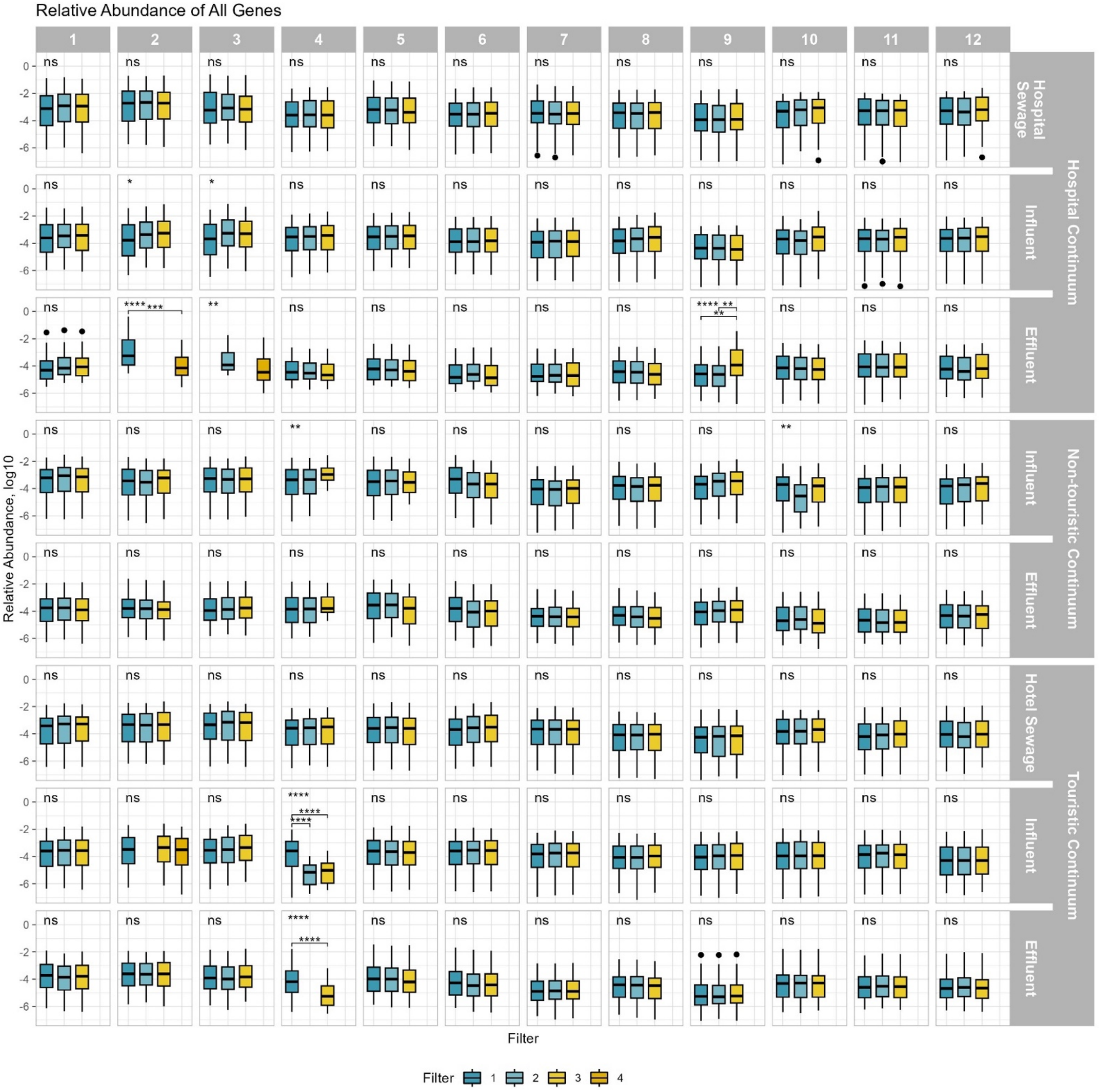


**Figure S3. Relative abundance of all ARGSs and MGEs before (influent) and after (effluent) wastewater treatment for all filters.** The horizontal numbers represent the sampling weeks.

Asterisks indicate significant differences between filters (* $p<0.05$, ** $p<0.01$, *** $p<0.001$, **** $p<0.0001$).

*Inter-Filter Variability*

In the hospital continuum, during the third campaign (week 9), Filter 3 exhibited a significantly higher relative abundance compared to the other two filters and was therefore removed from the analysis (Figure S3). *(Note: Data from Campaign 1 had already been excluded.)*

In the non-touristic continuum, statistical differences were observed between samples from weeks 4 and 10 of the influent sample (Figure S3).

In the touristic continuum, Filters 2 and 3 in week 4 (influent) displayed significantly lower relative abundance values than Filter 1. (Figure S3). While examining only week 4, Filters 2 and 3 appeared more similar to each other, suggesting that Filter 1 might be an outlier. However, when considering data from all filters across multiple time points, the overall trend indicated that Filters 2 and 3 were the likely outliers. Statistical differences were also observed between samples from week 4 of the effluent sample point.

Based on these results, the following technical replicates were removed:

- Hospital continuum, week 9 – Effluent sample, Filter 3, all replicates (Inter-filter variability)
- Touristic continuum, week 7 – Hotel sewage sample, Filter 3, Replicate 2 (Intra-filter variability)
- Touristic continuum, week 4 – Influent sample, Filters 2 and 3, all replicates (Intra-filter + Inter-filter variability)

### Tables

**Table S1**

Characteristics of the eight sampling points.

| Sampling point | Name | Description | Lat, long |
|---|---|---|---|
| 1 | Hospital sewage | Hospital sewer before the water joins the municipal water system | 16.2374, -61.5252 |
| 2 | Hospital: Influent | At the entrance of WWTP A (mixed hospital and municipal wastewater) | 16.2333, -61.55 |
| 3 | Hospital: Effluent | At the exit of WWTP A | 16.2333, -61.55 |
| 4 | Non-touristic: Influent | At the entrance of WWTP B (municipal wastewater from the small town and poultry farms) | 16.284596, -61.631024 |
| 5 | Non-touristic: Effluent | At the exit of WWTP B | 16.284596, -61.631024 |
| 6 | Hotel sewage | Wastewater coming from multiple hotels | 16.207970, -61.504663 |
| 7 | Touristic: Influent | At the entrance of WWTP C (mixed hotel and municipal wastewater) | 16.216156, -61.504035 |
| 8 | Touristic: Effluent | At the exit of WWTP C | 16.216156, -61.504035 |

**Table S2**

Resistome, exposome, and metadata collected at influent and effluent sampling points. Resistome: 80 targeted antibiotic-resistant genes (ARGs) grouped into 16 resistance classes and 12 mobile genetic elements (MGEs) including three integrons. Exposome: antibiotics, biocides, non-steroidal anti-inflammatory drugs, and heavy metals. Metadata: physicochemical properties and climate parameters. 16 clinically relevant ARGs and MGEs are in bold.

| | **Genes conferring resistance to:** | **ARGs and MGEs** |
|---|---|---|
| **RESISTOME** | Chloramphenicol | catB3, cat, cmlA1 |
| | Aminoglycosides | aac(3)-lid, aadE, **aac(6')-lb**, aadA, aac(6')-llA, spc, **aph(3')-III**, aph(2)-l(de), aph(2)-lb, aadB, AAC(3)-lb, strB, aac(6)-aph(2) |
| | Bacitracin | bacA_2, bacA_1 |
| | Beta-lactams | cepA, cblA, cfxA, **blaTEM**, blaIMP, blaAmpC, blaDHA, blaCMY-2, blaACC, **blaSHV**, **blaVIM**, **blaKPC**, blaIMI, blaBIC-1, blaGES, **blaNDM**, **blaOXA**, **blaCTX-M**, blaPER-1 |
| | Macrolides | ermF, **ermB**, ermC, ermY, mfsA, mefA_10, macB, ermX |

| | | |
|---|---|---|
| | Efflux pumps | acrA, mdtO, mdtL, mdtF, tolC |
| | Quinolones | qnrA, qnrB, **qnrS**, qnrC |
| | Heavy metals | cusF, copA, copD, cadA, merA, czcA |
| | QACs | qacA, qacC, qacE |
| | Vancomycin | vanA, vanB |
| | Tetracyclines | tetQ, tetW, **tetM**, tetO, tetB |
| | Polymyxin | arnA, **mcr-1** |
| | Sulfonamides | **sul1**, sulA |
| | Methicillin | **mecA** |
| | Trimethoprim | dfrA27, dfrF, dfrB1 |
| | Streptogramins | vat(A), vatB |
| | **Mobile genetic elements (MGEs):** | ISSW1, ISS1N, IS6_IS6100, Tp614, IS613, IS6 group, tn3_tnpA, ISEc9, inc-P1, integrons (**intI1**, intI2, intI3) |
| **EXPOSOME** | **Compound** | **Name** |
| | Antibiotic (ng/L) | Azithromycin, Ciprofloxacin, Erythromycin, Ofloxacine, Pyrimethamine, Sulfaguanidine, Sulfamethoxazole, Sulfathiazole, Trimethoprime |
| | Biocide (ng/L) | Acephate, Albendazole, Flubendazole, Levamisole, Pyrantel pamoate, Sulfoxaflor, Tebuconazole, Thiabendazole |
| | Anti-inflammatory (ng/L) | Diclofenac, Ibuprofen, Ketoprofen |
| | Heavy metal (µg/L) | As, Ba, Be, Bi, Cd, Ce, Co, Cr, Cu, Fe, Gd, Hg, Li, Mg, Mn, Mo, Pd, Pb, Pt, Rh, Sb, Se, Sn, Sr, Th, Tl, U, V, W, Zn |
| **METADATA** | **Type of data** | **Measurement** |
| | Physicochemical properties | Ammonium mg/L N,<br>BOD5 mg/O2/L,<br>TSS mg/L,<br>Nitrate mg/L N,<br>pH unit pH,<br>Total Phosphorus mg(P)/L,<br>COD mg(O2)/L,<br>Total nitrogen mg/L N<br><br>(BOD = biochemical oxygen demand; TSS = total suspended solids; COD = chemical oxygen demand) |
| | Climate parameters | Max temperature (°C),<br>Min temperature (°C),<br>Water temperature (°C),<br>Precipitation (mm) D-2,<br>Precipitation (mm) D-1,<br>Precipitation (mm) D0<br>(D-2 = two days before the sampling, D-1= day before the sampling, D0 = day of the sampling) |

**Table S3**

Average relative abundance of 16 genes of interest and the concentration of antibiotics (ng/L), biocides (ng/L), NSAIDs (ng/L), and heavy metals (μg/L) in influent and effluent wastewater samples across all campaigns. Values in red represent an increase in relative abundance or concentration after wastewater treatment. Statistically significant changes are represented in **bold**. Statistical significance was assessed using the non-parametric Wilcoxon signed-rank test for paired samples where p-values were adjusted using the Benjamini-Hochberg method.

| Gene of interest | Hospital continuum | | | Non-touristic continuum | | | Touristic continuum | | |
|---|---|---|---|---|---|---|---|---|---|
| | Influent | Effluent | % change | Influent | Effluent | % change | Influent | Effluent | % change |
| *aac(6')-lb* | 0.0036 | 0.0006 | **-83.0** | 0.0023 | 0.0068 | 200.7 | 0.0021 | 0.0006 | **-69.9** |
| *aph(3')-III* | 0.0001 | 2.05E-05 | **-84.9** | 0.0005 | 0.0001 | **-74.2** | 0.0002 | 6.82E-05 | **-69.7** |
| *blaCTX-M* | 3.58E-05 | 2.35E-05 | -34.3 | 1.25E-05 | 7.59E-06 | -39.3 | 2.03E-05 | 5.34E-06 | **-73.7** |
| *blaKPC* | 0.0002 | 8.79E-05 | -53.7 | 5.52E-05 | 4.06E-05 | -26.5 | 6.26E-05 | 2.22E-05 | **-64.5** |
| *blaNDM* | 7.76E-07 | 2.71E-07 | -65.1 | 6.08E-07 | 8.41E-07 | 38.3 | 9.24E-07 | 1.00E-08 | **-98.9** |
| *blaOXA* | 2.76E-05 | 8.78E-06 | **-68.2** | 5.94E-05 | 6.62E-06 | **-88.9** | 1.84E-05 | 3.40E-06 | **-81.5** |
| *blaSHV* | 1.53E-05 | 3.75E-06 | **-75.5** | 3.07E-05 | 1.10E-05 | **-64.0** | 2.01E-05 | 6.57E-06 | **-67.4** |
| *blaTEM* | 0.0019 | 0.0004 | **-79.6** | 0.0010 | 0.0004 | **-63.2** | 0.0010 | 0.0003 | **-72.5** |
| *blaVIM* | 4.27E-06 | 3.88E-06 | -9.0 | 3.74E-05 | 7.50E-06 | -80.0 | 8.42E-06 | 2.13E-06 | **-74.7** |
| *ermB* | 0.0028 | 0.0005 | **-80.7** | 0.0071 | 0.0011 | **-84.6** | 0.0070 | 0.0012 | **-82.6** |
| *intI1* | 0.0003 | 2.79E-05 | **-89.9** | 0.0007 | 0.0001 | **-85.2** | 0.0004 | 4.84E-05 | **-87.4** |
| *mcr-1* | 1.73E-07 | 1.65E-06 | **852.0** | 1.10E-06 | 4.76E-07 | -56.9 | 6.02E-08 | 1.00E-08 | -83.4 |
| *mecA* | 2.35E-05 | 1.84E-06 | **-92.2** | 3.91E-06 | 2.28E-06 | -41.6 | 1.93E-06 | 2.85E-06 | 48.0 |
| *qnrS* | 0.0047 | 0.0013 | **-72.7** | 0.0047 | 0.0019 | **-59.8** | 0.0028 | 0.0005 | **-81.5** |
| *sul1* | 0.0043 | 0.0012 | **-73.0** | 0.0040 | 0.0034 | -16.2 | 0.0028 | 0.0018 | **-34.5** |
| *tetM* | 0.0021 | 0.0004 | **-79.5** | 0.0021 | 0.0006 | **-72.3** | 0.0017 | 0.0005 | **-70.4** |
| **ANTIBIOTICS** | | | | | | | | | |
| AZ | 45.0 | 4.3 | -90.4 | 112.3 | 67.8 | -39.6 | 62.6 | 37.4 | -40.2 |
| CIP | 4.4 | 1.3 | -70.3 | 8.5 | 3.4 | -60.2 | 12.0 | 4.3 | **-64.4** |
| ERY | 67.7 | 17.8 | -73.7 | 145.8 | 104.1 | -28.6 | 90.5 | 86.3 | -4.6 |
| OFX | 10.5 | 5.8 | -45.1 | 7.3 | 11.2 | 53.5 | 22.5 | 13.8 | -38.8 |
| PM | 8.1 | 0.0 | -100.0 | 0.2 | 0.7 | 303.5 | 6.9 | 1.6 | -76.4 |
| SGN | 18.1 | 6.5 | -63.9 | 8.3 | 10.6 | 27.0 | 10.2 | 5.8 | -43.6 |
| SMX | 15.1 | 15.2 | 0.8 | 36.2 | 44.1 | 21.8 | 46.3 | 21.8 | -52.8 |
| STZ | 0.2 | 0.2 | -18.1 | 5.5 | 5.2 | -5.7 | 1.5 | 0.9 | -41.5 |
| TMP | 21.1 | 4.0 | -81.0 | 97.5 | 20.6 | **-78.8** | 67.5 | 19.7 | **-70.8** |
| **BIOCIDES** | | | | | | | | | |
| ACE | 46.2 | 17.4 | -62.4 | 88.7 | 51.2 | -42.3 | 40.2 | 38.3 | -4.7 |
| ABZ | 8.0 | 4.2 | -47.4 | 45.9 | 39.2 | -14.6 | 2.9 | 2.3 | -20.6 |
| FLU | 233.8 | 72.4 | -69.0 | 490.9 | 375.8 | -23.4 | 310.7 | 315.9 | 1.7 |

| | | | | | | | | | |
|---|---|---|---|---|---|---|---|---|---|
| LEV | 37.7 | 54.2 | 43.8 | 112.0 | 59.7 | -46.7 | 63.9 | 49.5 | -22.5 |
| PYR | 2.3 | 4.0 | 71.1 | 2.7 | 2.6 | -5.8 | 4.8 | 3.4 | -29.6 |
| SUL | 6.0 | 0.0 | -100.0 | 2.3 | 0.3 | -86.6 | 0.1 | 0.4 | 342.0 |
| TEB | 31.2 | 24.2 | -22.2 | 59.4 | 54.3 | -8.6 | 18.1 | 31.2 | 72.1 |
| TBZ | 7.3 | 7.2 | -1.1 | 20.9 | 12.2 | -41.4 | 15.3 | 20.4 | 33.2 |
| **NSAIDs** | | | | | | | | | |
| Diclofenac | 105.3 | 56.3 | -46.5 | 755.5 | 450.5 | -40.4 | 487.0 | 499.8 | 2.6 |
| Ibuprofen | 241.0 | 96.8 | -59.9 | 457.8 | 235.1 | -48.6 | 435.8 | 292.9 | -32.8 |
| Ketoprofen | 272.6 | 26.5 | **-90.3** | 335.6 | 130.6 | -61.1 | 293.4 | 216.6 | -26.2 |
| **HEAVY METALS** | | | | | | | | | |
| As | 3636.1 | 3316.4 | -8.8 | 1242.3 | 1051.8 | -15.3 | 5260.3 | 5233.8 | -0.5 |
| Cd | 29.3 | 33.9 | 15.7 | 36.3 | 24.5 | **-32.5** | 46.0 | 27.4 | -40.5 |
| Cr | 1344.5 | 941.7 | -30.0 | 1715.4 | 1422.7 | -17.1 | 2175.5 | 2167.6 | -0.4 |
| Cu | 8108.2 | 8899.9 | 9.8 | 12246.8 | 9666.7 | **-21.1** | 13334.6 | 10969.6 | -17.7 |
| Gd | 185.5 | 157.1 | -15.3 | 785.3 | 395.0 | -49.7 | 488.9 | 510.7 | 4.5 |
| Hg | 19.0 | 16.5 | -13.3 | 24.8 | 15.8 | **-36.2** | 22.9 | 16.6 | -27.7 |
| Pb | 1956.5 | 1886.5 | -3.6 | 2202.6 | 1883.1 | -14.5 | 3564.3 | 2411.9 | -32.3 |
| Zn | 33040.4 | 25662.2 | -22.3 | 58385.4 | 34307.4 | -41.2 | 60100.9 | 40596.8 | -32.5 |

#### Table S4

Evolution in the relative abundance of ARGs and MGEs of interest (normalized to 16S rRNA gene) and the concentration of antibiotics, biocides, NSAIDs, and heavy metals after WWTP. Statistical significance was assessed using the non-parametric Wilcoxon signed-rank test for paired samples. P-values in **bold** indicate a statistically significant change in effluent water samples.

*Benjamini-Hochberg adjustment method of p-values.

| ARG or MGE | HOSPITAL CONTINUUM | | NON-TOURISTIC CONTINUUM | | TOURISTIC CONTINUUM | |
|---|---|---|---|---|---|---|
| | p-value | Adj. p-value* | p-value | Adj. p-value* | p-value | Adj. p-value* |
| *aac(6')-lb* | 0.004 | **0.006** | 0.043 | 0.068 | <0.001 | **0.001** |
| *aph(3')-III* | 0.004 | **0.006** | <0.001 | **0.001** | <0.001 | **0.002** |
| *blaCTX-M* | 0.098 | 0.112 | 0.266 | 0.327 | <0.001 | **0.001** |
| *blaKPC* | 0.074 | 0.091 | 0.407 | 0.465 | 0.004 | **0.006** |
| *blaNDM* | 0.402 | 0.429 | 0.944 | 0.944 | 0.023 | **0.026** |
| *blaOXA* | 0.012 | **0.017** | <0.001 | **0.001** | 0.002 | **0.004** |
| *blaSHV* | 0.004 | **0.006** | <0.001 | **0.001** | <0.001 | **0.01** |
| *blaTEM* | 0.004 | **0.006** | <0.001 | **0.001** | <0.001 | **0.01** |
| *blaVIM* | 1 | 1 | 0.554 | 0.591 | <0.001 | **0.001** |
| *ermB* | 0.004 | **0.006** | <0.001 | **0.001** | <0.001 | **0.001** |

| | | | | | | |
|---|---|---|---|---|---|---|
| *intI1 (MGE)* | 0.004 | **0.006** | <0.001 | **0.002** | 0.006 | **0.008** |
| *mcr-1* | 0.036 | 0.048 | 0.058 | 0.084 | 0.371 | 0.396 |
| *mecA* | 0.004 | **0.006** | 0.034 | **0.061** | 0.541 | 0.541 |
| *qnrS* | 0.004 | **0.006** | <0.001 | **0.002** | <0.001 | **0.001** |
| *sul1* | 0.004 | **0.006** | 0.11 | 0.147 | 0.019 | **0.024** |
| *tetM* | 0.004 | **0.006** | <0.001 | **0.001** | <0.001 | **0.001** |
| **ANTIBIOTICS** | | | | | | |
| AZ | 0.012 | 0.053 | 0.064 | 0.173 | 0.339 | 0.509 |
| CIP | 0.012 | 0.053 | 0.012 | 0.055 | 0.004 | **0.031** |
| ERY | 0.074 | 0.134 | 0.077 | 0.173 | 1 | 1 |
| OFX | 0.129 | 0.166 | 0.38 | 0.545 | 0.037 | 0.096 |
| PM | 0.1 | 0.15 | 0.423 | 0.545 | 0.855 | 0.962 |
| SGN | 0.027 | 0.061 | 0.622 | 0.700 | 0.043 | 0.096 |
| SMX | 1 | 1 | 0.424 | 0.545 | 0.339 | 0.509 |
| STZ | 1 | 1 | 1 | 1 | 0.584 | 0.751 |
| TMP | 0.020 | 0.059 | 0.005 | **0.044** | 0.007 | **0.031** |
| **BIOCIDES** | | | | | | |
| ACE | 0.363 | 0.794 | 1 | 1 | 0.787 | 1 |
| ABZ | 1 | 1 | 0.424 | 1 | 0.308 | 0.493 |
| FLU | 0.020 | 0.156 | 0.824 | 1 | 0.91 | 1 |
| LEV | 0.496 | 0.794 | 0.092 | 0.738 | 0.266 | 0.493 |
| PYR | 0.441 | 0.794 | 0.906 | 1 | 0.236 | 0.493 |
| SUL | 0.371 | 0.794 | 1 | 1 | 1 | 1 |
| TEB | 0.82 | 0.937 | 0.91 | 1 | 0.129 | 0.493 |
| TBZ | 0.734 | 0.937 | 0.45 | 1 | 0.083 | 0.493 |
| **NSAIDs** | | | | | | |
| Diclofenac | 0.59 | 0.59 | 0.624 | 0.919 | 0.59 | 0.885 |
| Ibuprofen | 0.529 | 0.59 | 0.919 | 0.919 | 0.906 | 0.906 |
| Ketoprofen | 0.008 | **0.023** | 0.092 | 0.277 | 0.058 | 0.174 |
| **HEAVY METALS** | | | | | | |
| As | 0.563 | 0.651 | 0.055 | 0.086 | 0.82 | 0.937 |
| Cd | 1 | 1 | 0.004 | **0.016** | 0.131 | 0.32 |
| Cr | 0.031 | 0.250 | 0.413 | 0.413 | 0.16 | 0.32 |
| Cu | 0.094 | 0.375 | 0.004 | **0.016** | 0.027 | 0.218 |
| Gd | 0.57 | 0.651 | 0.266 | 0.304 | 0.97 | 0.97 |
| Hg | 0.219 | 0.438 | 0.012 | **0.031** | 0.074 | 0.297 |

| | | | | | | |
|---|---|---|---|---|---|---|
| Pb | 0.156 | 0.416 | 0.065 | 0.086 | 0.496 | 0.661 |
| Zn | 0.313 | 0.501 | 0.039 | 0.078 | 0.25 | 0.4 |

### Figures

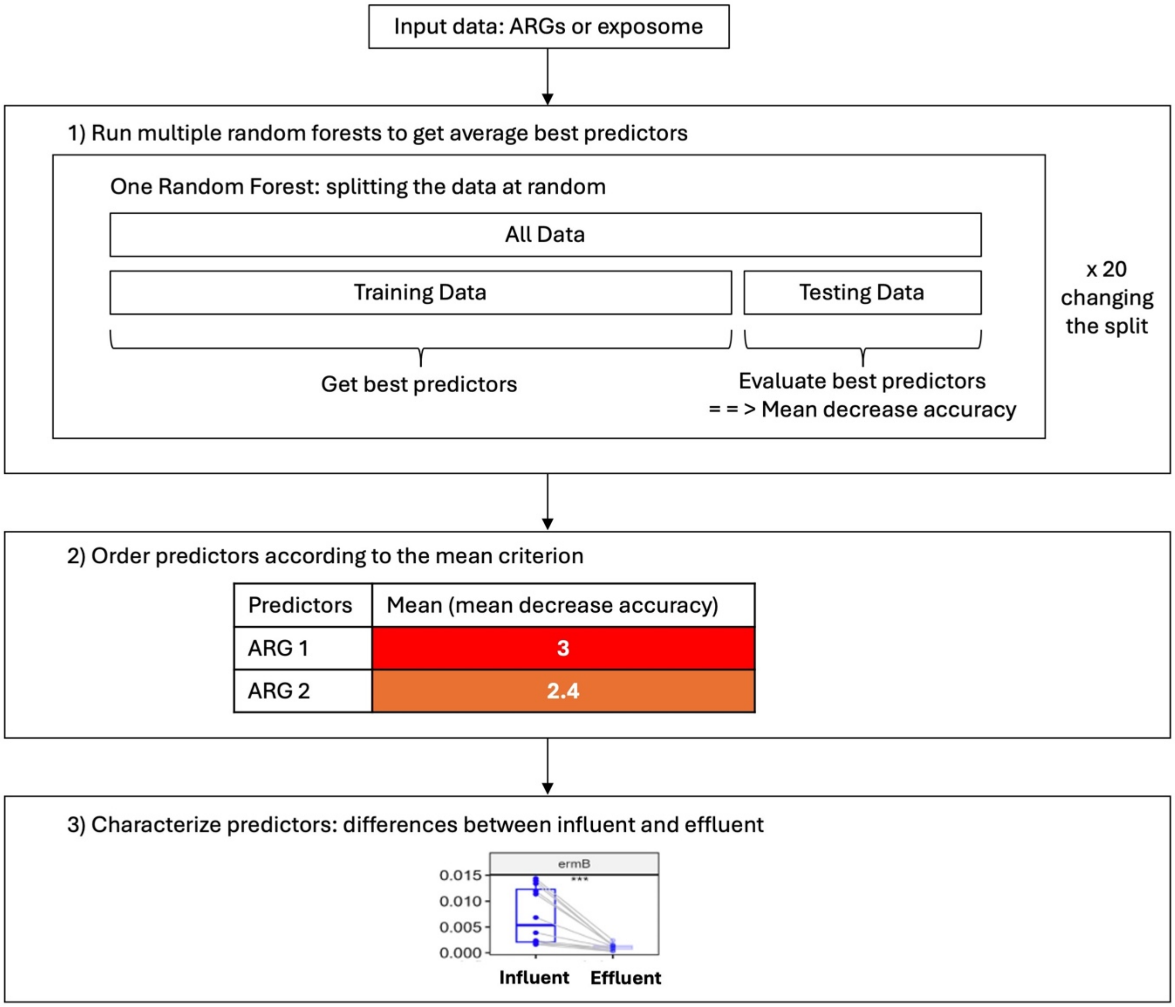


**Figure S4.** Schematic of the random forest analysis pipeline

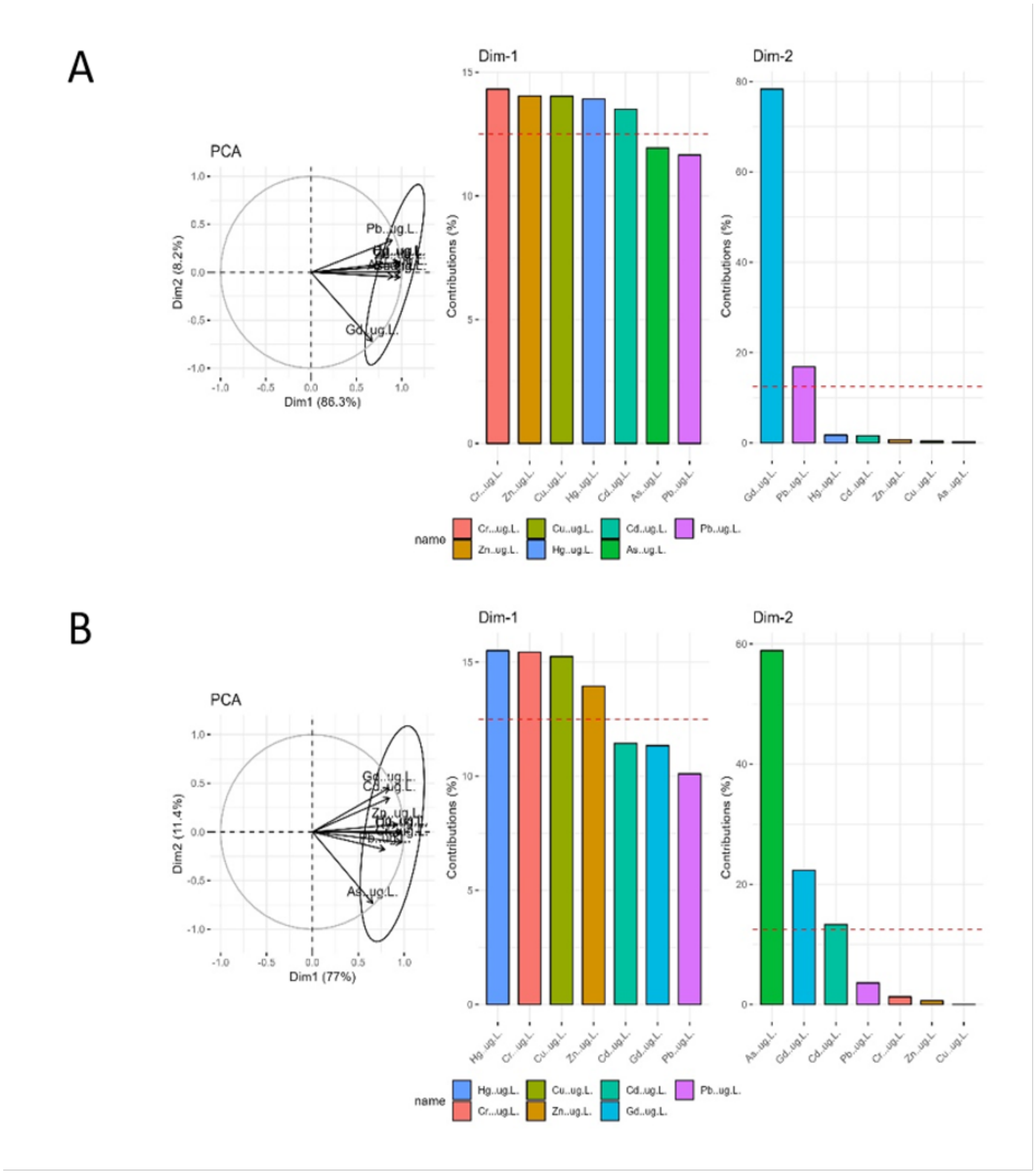


**Figure S5.** Principal component analysis (PCA) of heavy metals to identify and eliminate highly correlated variables within a hospital wastewater continuum. A. Influent B. Effluent

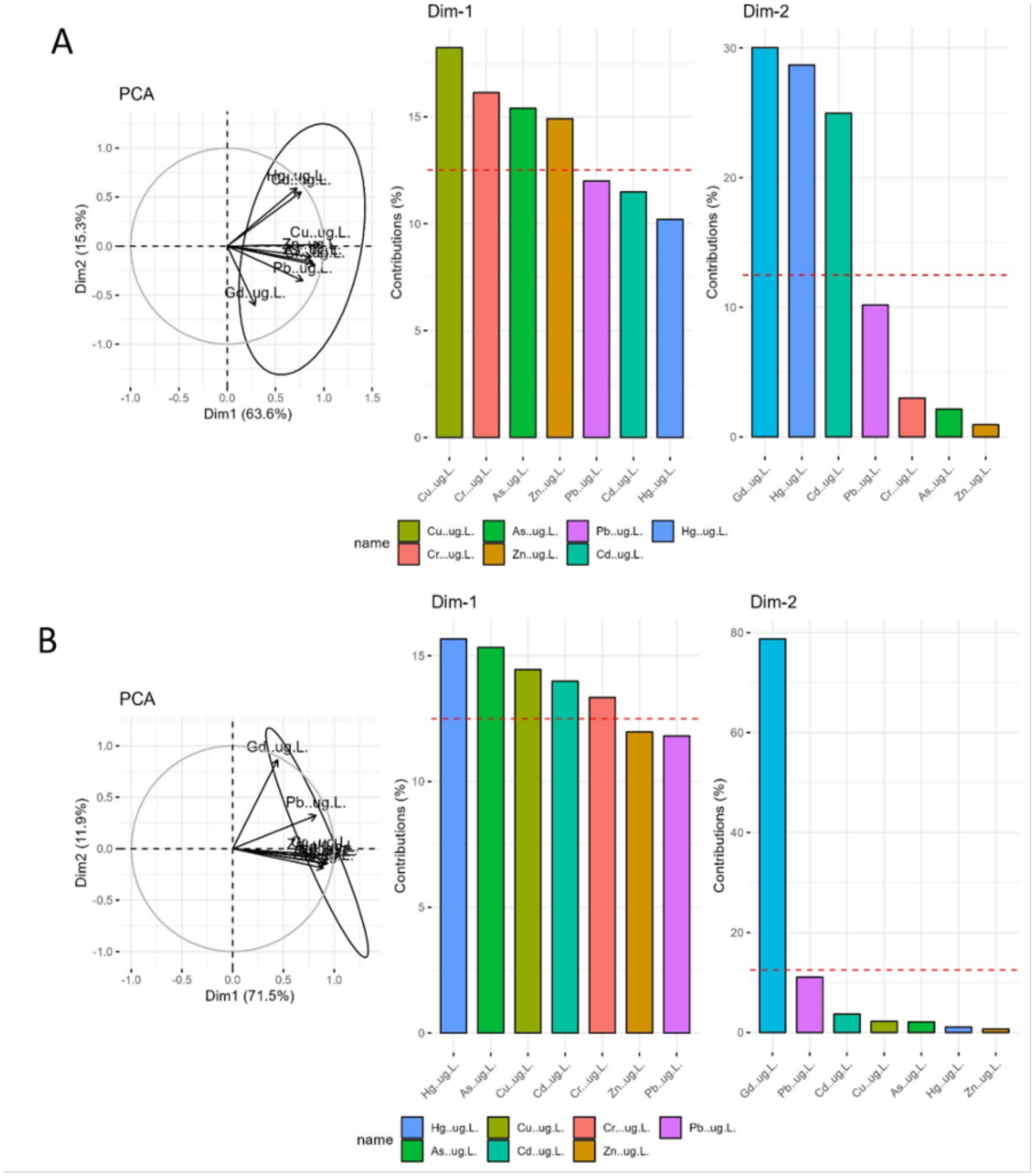


**Figure S6.** Principal component analysis (PCA) of heavy metals to identify and eliminate highly correlated variables within a non-touristic wastewater continuum. A. Influent B. Effluent

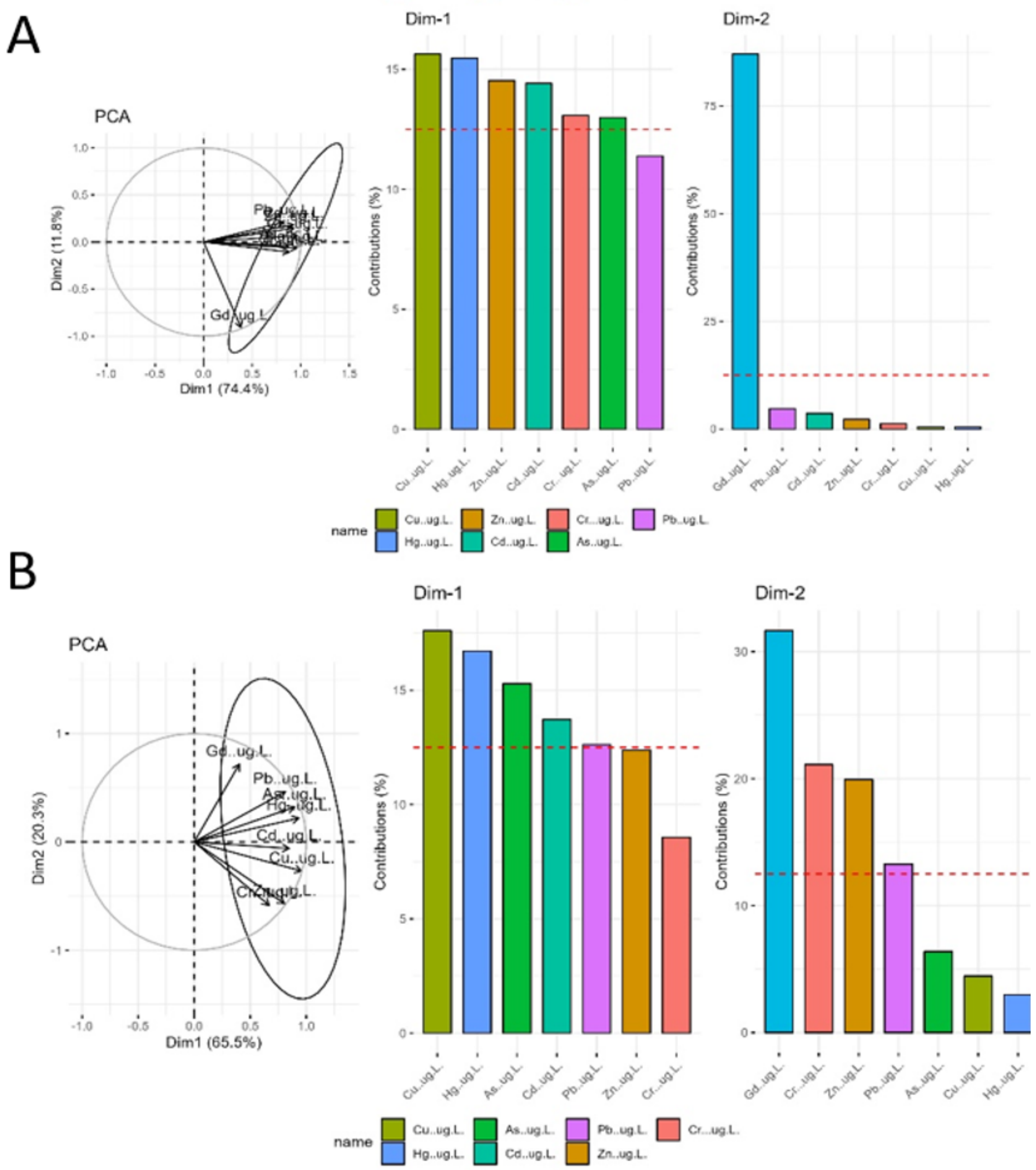


**Figure S7.** Principal component analysis (PCA) of heavy metals to identify and eliminate highly correlated variables within a touristic wastewater continuum. A. Influent B. Effluent

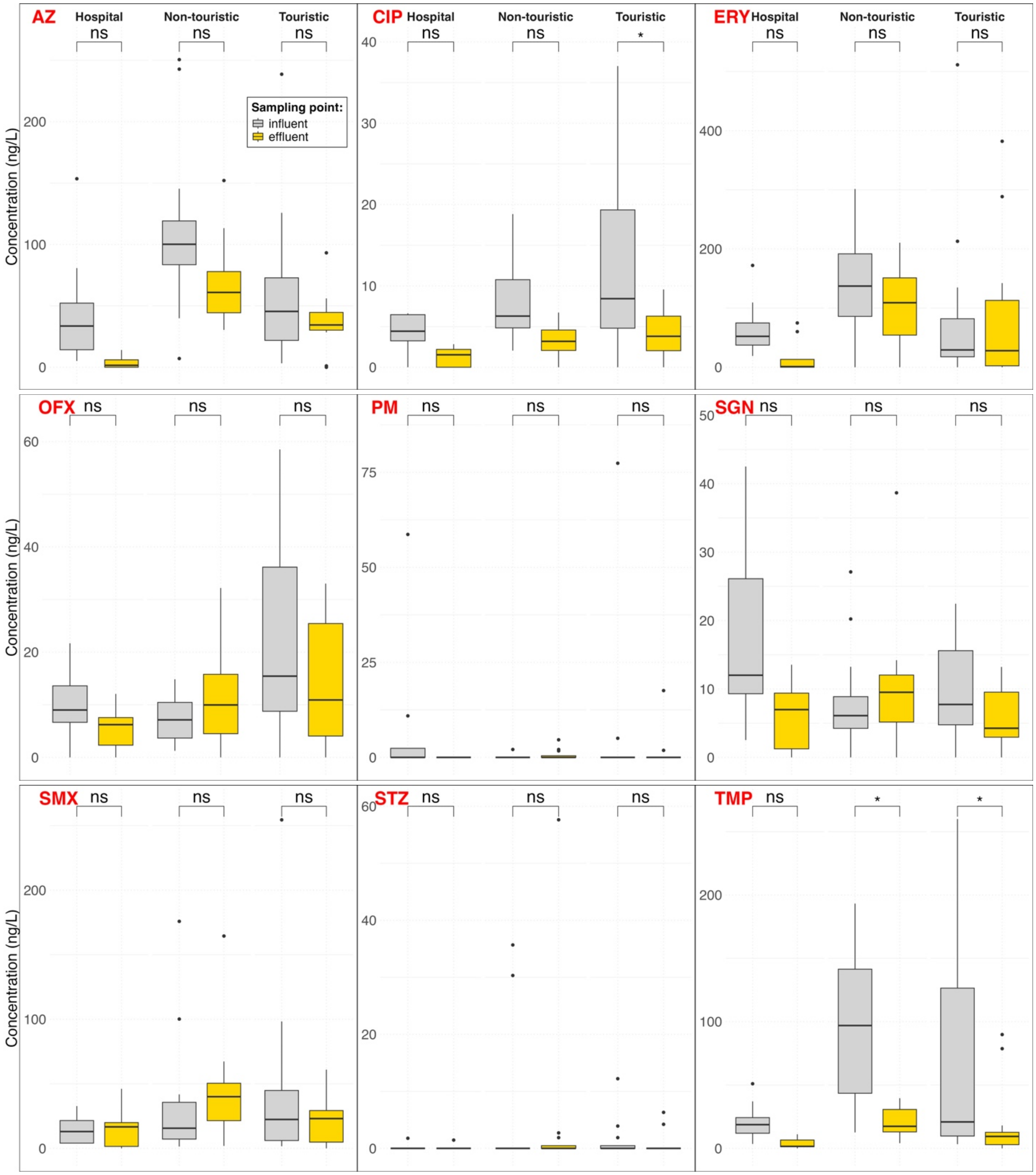


**Figure S8. Antibiotics.** Concentration (ng/L) of nine antibiotics in influent and effluent wastewater samples across hospital, non-touristic, and touristic wastewater continuums.

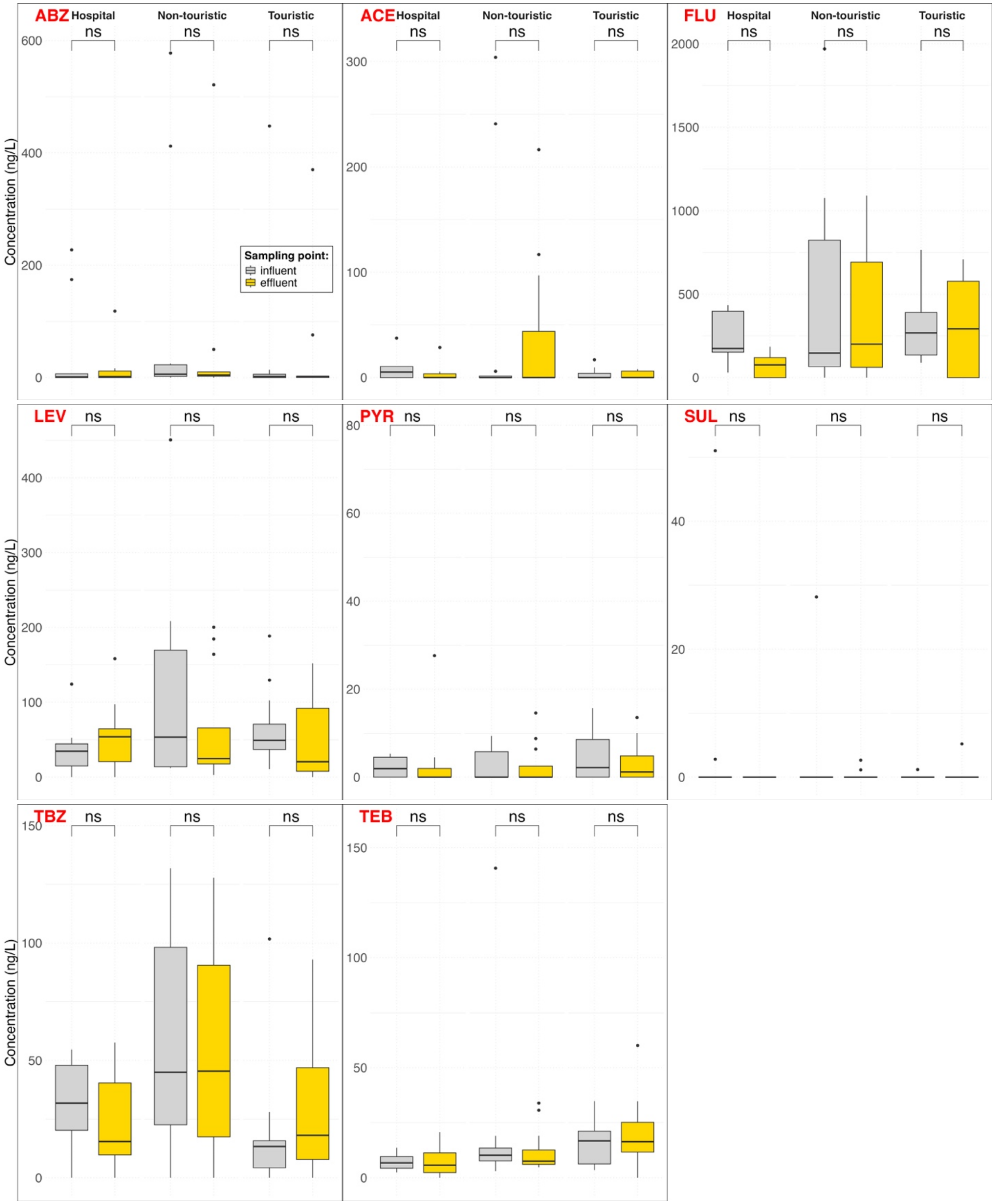


**Figure S9. Biocides.** Concentration (ng/L) of eight biocides in influent and effluent wastewater samples across hospital, non-touristic, and touristic wastewater continuums.

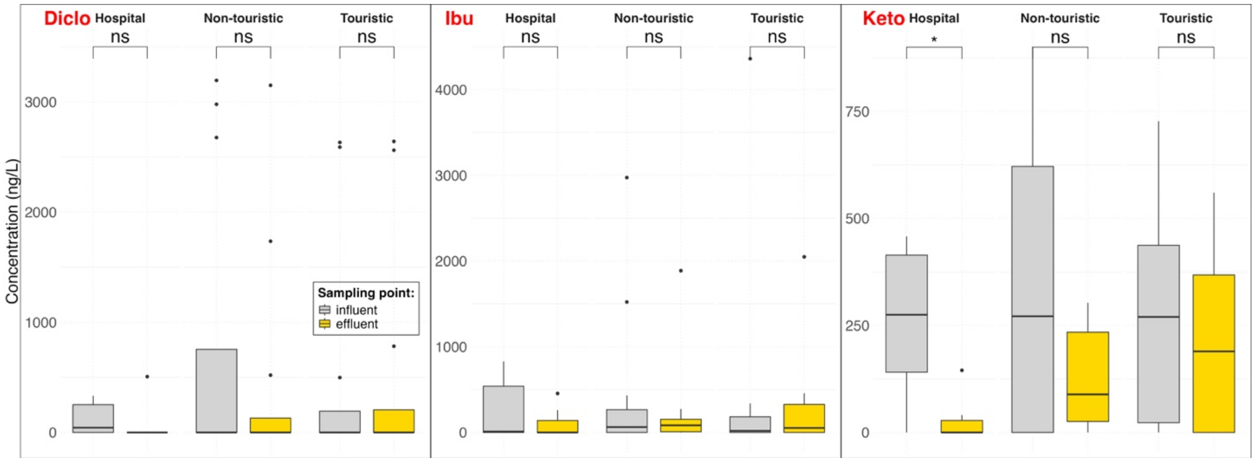


**Figure S10. Anti-inflammatory drugs.** Concentration (ng/L) of three non-steroidal anti-inflammatory drugs (diclofenac, ibuprofen, and ketoprofen) in influent and effluent wastewater samples across hospital, non-touristic, and touristic wastewater continuums.

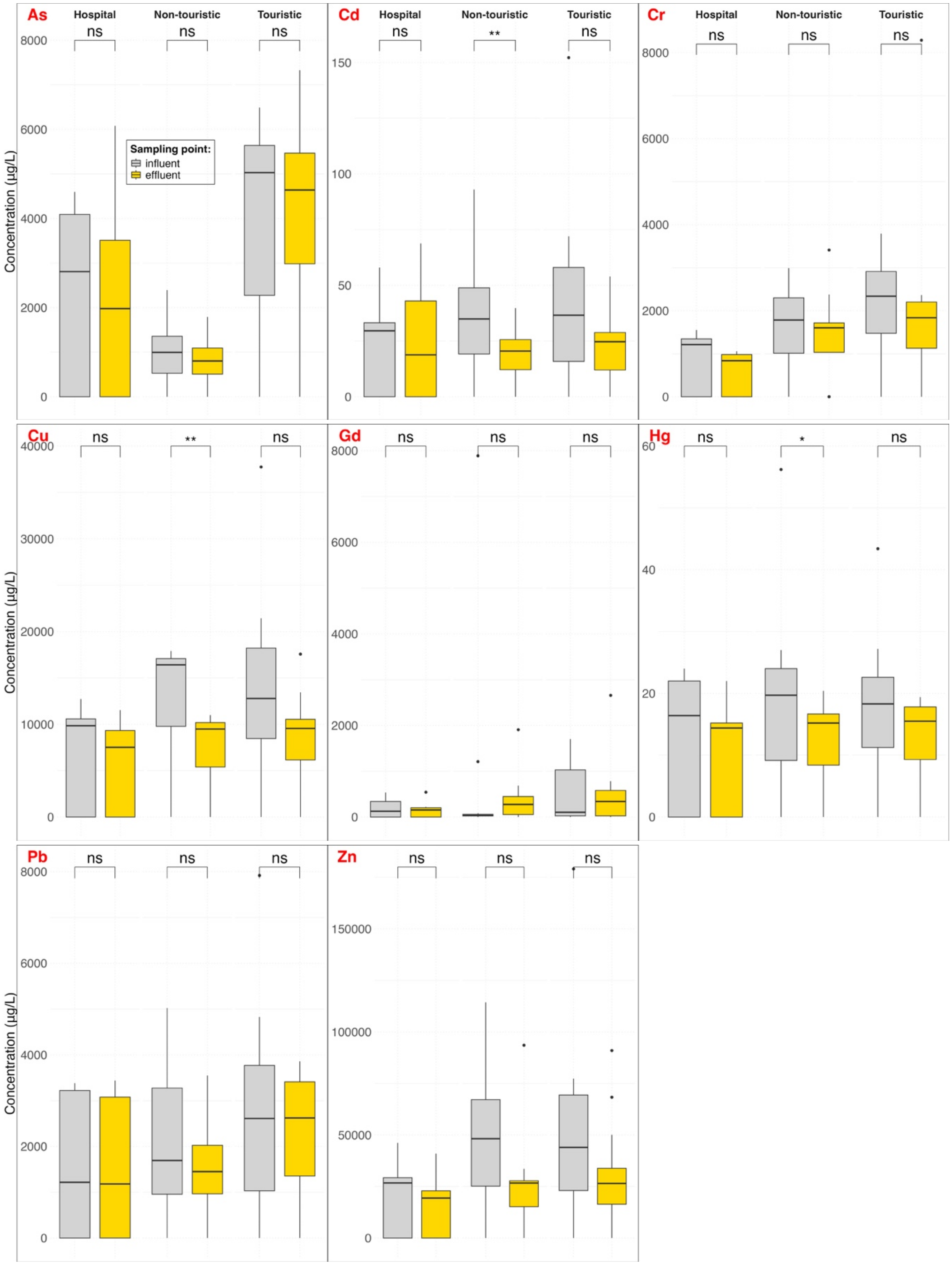


**Figure S11. Heavy metals.** Concentration (µg/L) of eight heavy metals in influent and effluent wastewater samples across hospital, non-touristic, and touristic wastewater continuums.
